# Fbxo42 promotes the degradation of Ataxin-2 granules to trigger terminal Xbp1 signaling

**DOI:** 10.1101/2024.12.22.629979

**Authors:** Cristiana C Santos, Nadine Schweizer, Fátima Cairrão, Juanma Ramirez, Nerea Osinalde, Ming Yang, Catarina J Gaspar, Vanya I Rasheva, Zach Hensel, Colin Adrain, Tiago N Cordeiro, Franka Voigt, Paulo A Gameiro, Ugo Mayor, Pedro M Domingos

**Affiliations:** Instituto de Tecnologia Química e Biológica António Xavier, Universidade Nova de Lisboa, Oeiras, Portugal; Department of Biochemistry and Molecular Biology, University of the Basque Country (UPV/EHU), Leioa, Bizkaia, Spain; Department of Molecular Life Sciences, University of Zurich, Switzerland; Patrick G Johnston Centre for Cancer Research, Queen’s University, Belfast, UK; NOVA Medical School Faculdade de Ciências Médicas, NMS FCM, Universidade Nova de Lisboa, Lisboa, Portugal; Ikerbasque, Basque Foundation for Science, Bilbao, Spain

## Abstract

The Unfolded Protein Response (UPR) is composed by homeostatic signaling pathways that are activated by the accumulation of misfolded proteins in the Endoplasmic Reticulum (ER), a condition known as ER stress. Prolonged ER stress and activation of the UPR causes cell death, by mechanisms that remain poorly understood. Here, we report that regulation of Ataxin-2 by Fbxo42 is a crucial step during UPR-induced cell death. From a genetic screen in *Drosophila*, we identified loss of function mutations in *Fbxo42* that suppress cell death and retinal degeneration induced by the overexpression of Xbp1^spliced^, an important mediator of the UPR. We identified the RNA binding protein Ataxin-2 as a substrate of Fbxo42, which, as part of a Skp-A/Cullin-1 complex, promotes the ubiquitylation and degradation of Ataxin-2. Upon ER-stress, the mRNA of Xbp1 is not immediately translated but instead it is sequestered and stabilized in Ataxin-2 granules, until the Fbxo42 recruitment to these granules promotes the degradation of Ataxin-2, allowing the translation of Xbp1 mRNA at later stages of UPR activation. Our results identify Fbxo42-mediated degradation of Ataxin-2 as a key mechanism regulating the levels of Xbp1 mRNA and protein, to trigger cell death during the terminal stages of UPR activation.

## INTRODUCTION

The Endoplasmic Reticulum (ER) is the cell organelle where secretory and membrane proteins are synthesized and folded. When the folding capacity of the ER is impaired, the presence of unfolded or incorrectly folded (misfolded) proteins in the ER causes stress on the cell - ER stress - and activates the Unfolded Protein Response (UPR), to restore homeostasis in the ER (Walter and Ron, 2011).

The UPR is accomplished via the activation of signaling pathways induced by 3 ER transmembrane molecular sensors that detect ER stress: Inositol-requiring enzyme 1 (Ire1), Pancreatic ER kinase (PKR)-like ER kinase (Perk) and Activating transcription factor 6 (Atf6). The activation of the UPR leads to the reduction of general protein synthesis, to prevent further accumulation of proteins in the ER, and the transcriptional up-regulation of a specific subset of genes, including genes encoding ER chaperones and enzymes, to increase the protein folding capacity of the ER. However, when ER stress is prolonged and severe, leading to chronic activation of the UPR, cells can activate apoptosis, a genetic programmed form of cell death, by mechanisms that are still poorly understood (Rasheva and Domingos, 2009; Tabas and Ron, 2011).

The transcription factor Xbp1 is an important mediator of Ire1 signaling. Upon the activation of the UPR, the endoribonuclease domain of Ire1 cleaves the Xbp1 mRNA in two sites, a non-conventional splicing event that causes a frameshift during translation, introducing a longer carboxyl domain in the protein encoded by Xbp1^spliced^ (Xbp1s) mRNA (Walter and Ron, 2011). Only Xbp1^spliced^ is fully active as a transcription factor that regulates the expression of ER chaperones and other target genes (Acostaalvear et al., 2007).

Here, we find that persistent expression of Xbp1^spliced^ (Xbp1s) induces cell death, generating a smaller, atrophic “glossy” eye phenotype that is suppressed by co-expression of anti-apoptotic proteins. Through genetic screening, we identified loss of function mutations in the F-box protein Fbxo42 that suppress cell death induced by Xbp1^spliced^. Next, we performed a proteomic screen using pulldowns with a biotinylable form of ubiquitin (^bio^Ub) to identify the relevant substrates of Fbxo42 and discovered the RNA-binding protein Ataxin-2, as a key target. In cells undergoing high levels of ER stress, Xbp1 mRNA is initially stored in cytoplasmic Ataxin-2 granules, preventing its translation, until Fbxo42 is recruited to promote the degradation of Ataxin-2 granules, allowing for the translation of Xbp1 mRNA at terminal stages of UPR activation.

## RESULTS

### Expression of Xbp1^spliced^ causes an atrophic “glossy” eye phenotype

In *Drosophila*, loss of function mutations in Xbp1 increase photoreceptor degeneration caused by *ninaE* (rhodopsin-1) misfolding mutations (Ryoo et al., 2007). Surprisingly, in comparison with a “wild type” *Drosophila* adult eye (Fig.1A), expression of Xbp1^spliced^ under the control of GMR-GAL4 causes a smaller, atrophic “glossy” eye phenotype (Fig. 1B), where the adult *Drosophila* eye becomes a thin layer of yellow cuticle due to the lack of photoreceptors, cone and the red pigment cells (Liao et al., 2006). This “glossy” eye phenotype is suppressed by co-expression of Xbp1^unspliced^ (Xbp1u, Fig. 1C), P35 (Fig. 1D) or DIAP1 (Fig. 1E), with the partial recovery of the red pigment and the external lens structure. Partial suppression by Xbp1^unspliced^ is consistent with the role of mammalian Xbp1^unspliced^ in promoting the down-regulation of Xbp1^spliced^ signaling (Yoshida et al., 2009). Partial suppression by baculovirus P35 or *Drosophila* inhibitor of apoptosis 1 (DIAP1), two anti-apoptotic proteins, indicates that apoptotic cell death plays a role in the induction of the “glossy” eye phenotype by Xbp1^spliced^.

**Figure 1.**
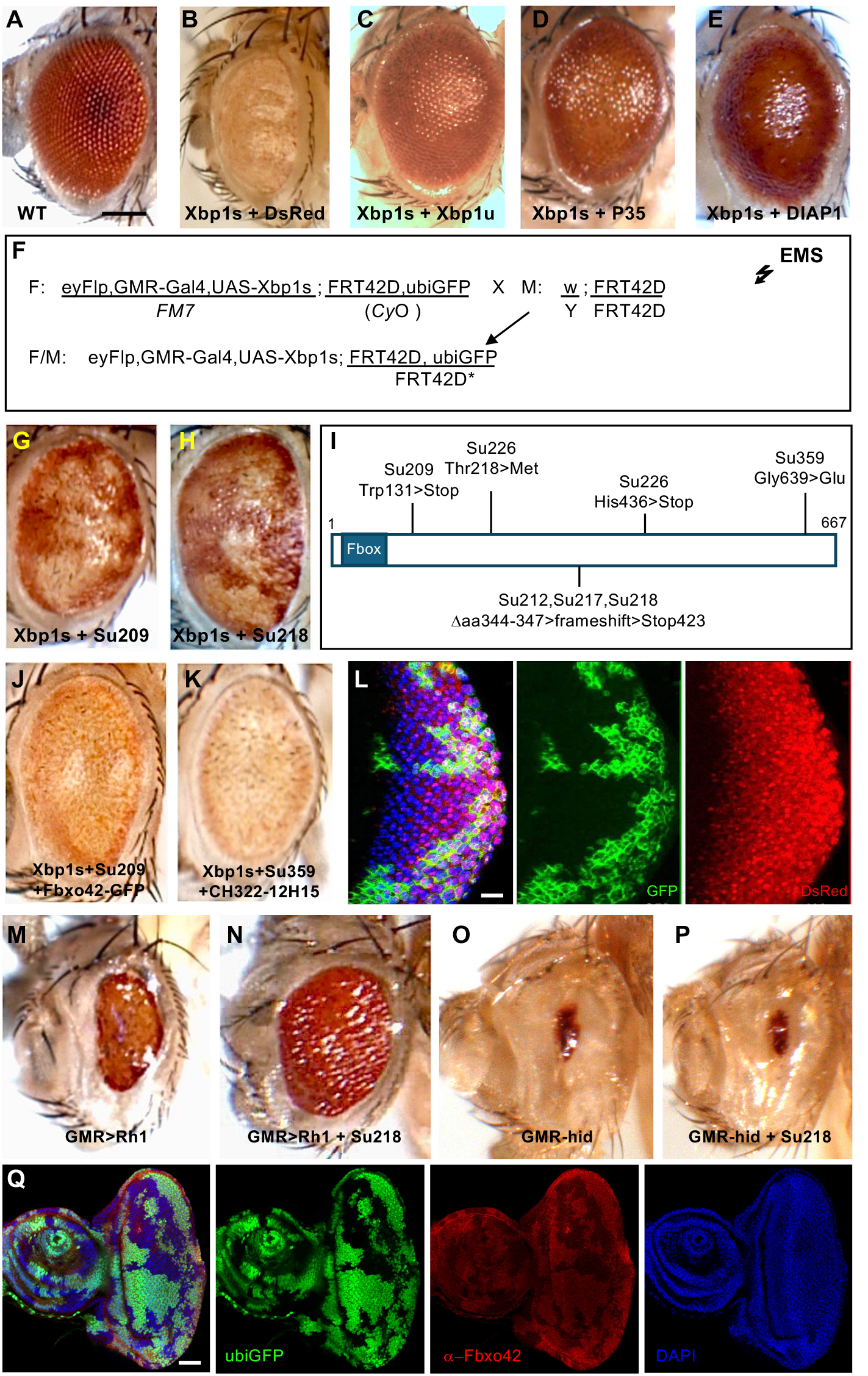
Loss of function mutations in Fbxo42 suppress glossy eye phenotype caused by overexpression of Xbp1^spliced^. A) Wild-type adult *Drosophila* eye (Canton S). Posterior is to the right and dorsal to the top in this and all subsequent panels. Scale bar = 200 μm. B) Glossy eye phenotype caused by overexpression of Xbp1^spliced^ (Xbp1s). Genotype: GMR-GAL4, UAS-Xbp1s, UAS-DsRed. C) Glossy eye phenotype is suppressed by co-expression of Xbp1^unspliced^ (Xbp1u) Genotype: GMR-GAL4, UASXbp1s, UAS-Xbp1u. D) Glossy eye phenotype is suppressed by co-expression of caspase inhibitor P35. Genotype: GMR-GAL4, UAS-Xbp1s, UAS-P35. E) Glossy eye phenotype is suppressed by co-expression of DIAP1. Genotype: GMR-GAL4, UAS-Xbp1s, UAS-DIAP1. F) Genetic scheme of the FLP/FRT F1 genetic screen for suppressors of the glossy eye phenotype caused by overexpression of Xbp1s. G) Glossy eye phenotype is suppressed by clones of suppressor 209. Genotype: eyFlp, GMR-GAL4, UAS-Xbp1s; FRT42D, Su209/FRT42D, ubiGFP. H) Glossy eye phenotype is suppressed by clones of suppressor 218. Genotype: eyFlp, GMR-GAL4, UAS-Xbp1s; FRT42D, Su218/FRT42D, ubiGFP. I) Schematic representation of Fbxo42 (amino acids 1 to 667), with the Fbox domain (blue box) and the mutations found in six of the suppressor alleles (Su209, Su212, Su217, Su218, Su226 and Su359) obtained from the genetic screen (f). Su212, Su217 and Su218 have a small deletion that causes a frameshift and premature Stop codon at 423 J) Overexpression of Fbxo42-GFP abolishes suppression of glossy eye by Su209. Genotype: eyFLP,GMR-GAL4; FRT42D, Su209/FRT42D, ubiGFP; UAS-Fbxo42-GFP. K) A genomic rescue construct containing Fbxo42 (Pacman CH322-12H15) abolishes suppression of glossy eye by Su359. Genotype: eyFLP,GMR-GAL4; FRT42D, Su359/FRT42D, ubiGFP; CH322-12H15. L) Immunofluorescence of 3^rd^ instar larva eye discs containing clones of Su218, labelled by the absence of ubiGFP (green). GMR-GAL4 driven expression of UAS-DsRed is similar in Su 218 and ubiGFP clones. Genotype: eyFlp,GMR-GAL4, UAS-DsRed; FRT42D, Su218/FRT42D, ubiGFP. Scale bar = 30 μm. M) Glossy eye phenotype caused by overexpression of Rh1. Genotype: eyFlp,GMR-GAL4; FRT42D/FRT42D, GMRhid, CL ;UAS-Rh1. N) Whole eye mutant clones of Su218 suppress glossy eye phenotype caused by overexpression of Rh1. Genotype: eyFlp,GMR-GAL4; FRT42D, Su218/FRT42D, GMRhid, CL ;UAS-Rh1. O) Small eye phenotype caused by overexpression of hid. Genotype: eyFlp,GMR-hid; FRT42D/FRT42D,ubiGFP. P) Clones of Su218 do not suppress the small eye phenotype caused by overexpression of hid. Genotype: eyFlp,GMR-hid; FRT42D, Su218/FRT42D,ubiGFP. Q) Immunofluorescence of 3^rd^ instar larva eye discs containing clones of Su218, labelled by the absence of ubiGFP (green), with an antibody made against Fbxo42 (Red). DAPI is in blue. Genotype: eyFlp,GMR-GAL4; FRT42D, Su218/FRT42D, ubiGFP. Scale bar = 60 μm.

Mosaic expression of a control UAS-DsRed (Fig. S1A), by the use of the Flipase/FRT technique (Golic, 1991), results in an adult eye where around two thirds of the cells have DsRed expression. Mosaic expression using a FRT chromosome containing both UAS-Xbp1^spliced^ and UAS-DsRed transgenes (Fig. S1B), results in a much reduced number of DsRed positive cells that present the “glossy” phenotype. These results indicate that Xbp1^spliced^ induces the “glossy” phenotype in a cell autonomous manner and that most of the cells expressing Xbp1^spliced^ die during eye development. In fact, higher levels of apoptotic TUNEL staining are observed in 3^rd^ instar larva eye discs containing clones of overexpression of Xbp1^spliced^, labelled by DsRed (Fig. S1C). We have also observed upregulation of the autophagy marker LC3-GFP by mosaic overexpression of Xbp1^spliced^ in the larval eye disc (Fig. S1D), consistent with the results previously reported for overexpression of Xbp1^spliced^ in the fat body (Arsham and Neufeld, 2009).

### Loss of function mutations in Fbxo42 suppress Xbp1^spliced^-induced “glossy” eye phenotype

In order to investigate the downstream molecular mechanisms involved in the induction of the “glossy” eye phenotype by Xbp1^spliced^, we performed a F1 genetic screen where mosaic eyes were generated by the Flipase/FRT technique (Golic, 1991) with Ethyl methanesulfonate (EMS) mutagenized chromosomes (Fig. 1F), in the background of Xbp1^spliced^ overexpression. Around 80.000 EMS mutagenized flies were screened, leading to the identification of 32 mutations in the right arm of the second chromosome (2R) that suppress the Xbp1^spliced^ “glossy” eye phenotype.

All the mutations were crossed to each other to create complementation groups by lethality. The biggest complementation group contained 14 mutant alleles, including suppressor (Su) 209 (Fig. 1G) and Su218 (Fig. 1H), which show clones with suppression of the “glossy” eye phenotype, with partial recovery of the red pigment and the external lens morphology in the eye. This complementation group was mapped by crossing the mutant alleles with a set of deficiencies (genomic deletions) covering 2R. From these crosses, we failed to obtain viable non-balanced progeny (except a few adult escapers with out-held wing phenotype) from Df(2R)BSC597, but we obtained viable non-balanced progeny from Df(2R)Exel7170 and Df(2R)01D01W-L133, mapping this complementation group to an interval containing 12 genes, between 58C1 and 58D1 (Fig. S1E). Viable non-balanced progeny was obtained from crosses to Df(2R)a(7) but not from Df(2R)a(EX1). Since the distal end of Df(2R)a(EX1) is not molecularly defined (Liu and Lengyel, 2000), we tested how far distally this deficiency reaches by crossing Df(2R)a(EX1) to lethal P element insertions in *Vps35* (P(EP)Vps35^EY16641^ and P(EP)Vps35^EY14200^) and *CG3074* (P(GawB)CG3074^NP7371^). Since viable non-balanced progeny was obtained from these crosses, we concluded that Df(2R)a(EX1) does not reach *Vps35*. Also, our suppressor mutations were not lethal over the lethal P element insertions in *Vps35*. With this approach, we rendered our region of interest to six genes (*Fbxo42, CG3045, CG11170, CG30279, CG11275* and *MED16*). Since no lethal alleles existed for these genes, we sequenced genomic DNA extracted from homozygous larvae for Su218, Su359 and the isogenized FRT42D line that was subjected to EMS mutagenesis. Our first attempt was with the gene *MED16*, in which no mutations were found, but in our second attempt we found several mutations in *Fbxo42* (*CG6758*). In total, we sequenced six suppressor alleles and all had mutations in *Fbxo42*, mostly premature STOP codons (Fig. 1I).

Mosaic suppression of the “glossy” eye phenotype by Fbxo42 mutants could be rescued by overexpression of Fbxo42-GFP (Fig. 1J) or by using a Pacman genomic rescue construct (CH322-12H15) containing Fbxo42 (Fig. 1K), demonstrating that the phenotype is Fbxo42 specific. We tested UAS-DsRed expression levels in clones of Fbxo42 mutants (Su218 - Fbxo42^N423*^) but saw no difference between the homozygous mutant and control tissue (Fig. 1L), which indicates that Fbxo42 is not required for transcriptional activation by GAL4/UAS. Su218 suppressed the strong rough eye phenotype induced by overexpression of rhodopsin-1 (Rh1) under the control of GMR-GAL4 (Fig. 1M,N), a paradigm where misfolded Rh1 accumulates in the ER, causing ER stress, activation of the UPR and Ire1-mediated splicing of Xbp1 (Kang and Ryoo, 2009)(Kang et al., 2012). However, Su218 did not suppress the small eye phenotype induced by overexpression of the proapoptotic gene *hid* (Fig. 1O,P), indicating that Fbxo42 acts downstream of UPR activation but upstream of the apoptotic machinery.

The genome of *Drosophila* has 45 F-box proteins (Dui et al., 2012) that form complexes with SkpA, Cullin-1, Roc1/Rbx1 and E2 ubiquitin ligases to mediate the ubiquitylation of specific substrates. In such complexes, F-box proteins bind to SkpA through the conserved F-box domain, which is present N-terminally in Fbxo42 (Fig. 1I). In *Drosophila* S2 cells (Fig. S1F), SkpA co-immunoprecipitates with FLAG-HA-Fbxo42, but not with a deletion of the F-box domain (FLAG-HA-ΔFbxo42), in accordance with results showing binding of Fbxo42 with SkpA in the *Drosophila* ovary (Barbosa et al., 2020).

We generated a rabbit polyclonal antibody raised against amino acids 384-667 of Fbxo42, which was used in immunofluorescence experiments with 3^rd^ instar larval eye discs containing clones of Su218 (Fbxo42^N423*^). Cells homozygous for Su218, labelled by the absence of ubiGFP, present reduced immunoreactivity for anti-Fbxo42, in comparison with neighboring control heterozygous cells (Fig. 1Q). In immunoblots, anti-Fbxo42 detects a band just above the 72 kDa marker that is absent in larva homozygous for Su226 (Fbxo42^H436*^, Fig. S1G). The expected size for Fbxo42 is 74,4 kDa. This specific band is present in several 3^rd^ instar larval tissues (Fig. S1H) and in the adult brain, testis and ovaries (Fig. S1K).

### Fbxo42 promotes the formation of Ataxin-2-ubiquitin conjugates

In order to identify the specific substrate(s) of Fbxo42, we used a proteomic approach, based on the ^bio^Ub strategy (Ramirez et al., 2015a). Transgenic flies with full-length FLAG-HA-Fbxo42, FLAG-HA-ΔFbxo42 or FLAG-Fblx7 (Bosch et al., 2014), a different F-box protein used as control, were crossed with ^bio^Ub flies (Ramirez et al., 2015a), a strain containing ubiquitin (Ub) with a short biotinylatable motif and BirA (biotin ligase), and GMR-GAL4, to drive the expression of the constructs in the *Drosophila* eye. We performed total protein extraction from FLAG-HA-Fbxo42, FLAG-HA-ΔFbxo42 and FLAG-Fblx7-expressing adult fly heads and confirmed expression of the F-box constructs and biotinylation of target proteins, by doing immunoblot with anti-FLAG and anti-Biotin antibodies (Fig. S2A). To purify the ubiquitylated proteins, we performed pulldowns using a high-capacity streptavidin resin from triplicate samples, which were then processed for identification by mass spectrometry (MS)(Fig. 2A, S2B and Table S1). The values observed for the endogenously biotinylated acetyl-coenzyme A and Pyruvate carboxylases (ACC and PCB) as well as for the overexpressed F-box proteins Fbxo42 and Fbxl7 (Fig. 2A) confirmed respectively that proportional amounts of biological material were isolated, and the overexpression of the corresponding F-box protein. In order to obtain results with statistical significance, stringent thresholds were established (Ramirez et al., 2015a)(Martinez et al., 2017), to minimize the number of false positives. This approach led to the identification of Ataxin-2 as a Fbxo42 substrate, since Ataxin-2 is more ubiquitylated upon Fbxo42 overexpression, in comparison with Fbxl7 overexpression (Fig. 2A and Table S1). Ataxin-2 is a RNA binding protein and participates in the formation of ribonucleoprotein granules (Ripin and Parker, 2023).

**Figure 2.**
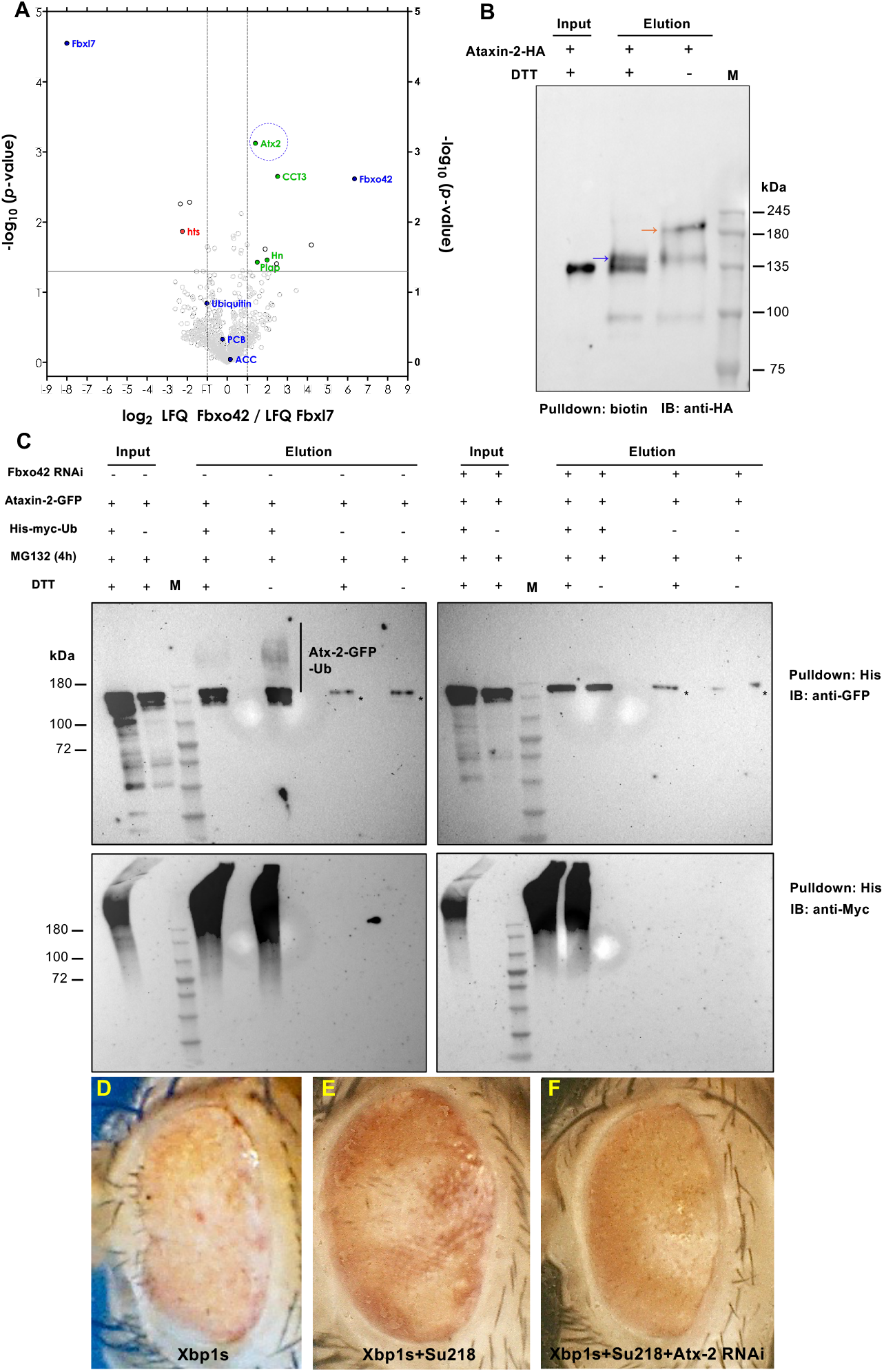
Fbxo42 promotes Ataxin-2/Ubiquitin conjugates in *Drosophila* eyes and S2 cells. A) Volcano plot of proteins identified by mass spectrometry after streptavidin/biotin pulldowns from *Drosophila* adult heads expressing ^bio^Ub (ubiquitin with biotinylation acceptor site) and Fbxo42 or Fbxl7, under the control of GMR-GAL4 driver. Results are presented as log2 LFQ (label free quantitation) intensity ratios. As blue dots are ACC and PCB, endogenous carboxylases which are biotinylated naturally and should have a log2 ratio of around 0, if similar amount of biological samples have been isolated during the pulldown. Also as blue dots are Ubiquitin, Fbxl7 and Fbxo42. As green and red dots are proteins identified by at least 2 unique peptides, which are either enriched or depleted, respectively. Ataxin-2 (Atx2) is highlighted by a blue dashed circle. Statistical analysis was performed by two-sided Student’s t test. The full results are provided in table S1. B) Immunoblot from *Drosophila* adult heads expressing ^bio^Ub, Fbxo42 and Ataxin-2-HA, under the control of GMR-GAL4 driver. *Drosophila* head lysates were subjected to streptavidin/biotin pulldowns and immunoblot with anti-HA. DTT sensitive Ataxin-2-HA bioUb conjugates are detected by an upwards shift of the bands on the gel. C) Immunoblot of protein extracts from *Drosophila* S2 cells expressing His-myc-Ub, Ataxin-2-GFP and in the presence or absence of RNAi against Fbxo42. S2 cells lysates were subjected to Histidine (His) pulldown and immunoblots with anti-GFP (top panels) or anti-myc (bottom panels). DTT sensitive Ataxin-2-GFP His-mycUb conjugates are detected by an upwards shift of the bands on the gel and are not observed upon Fbxo42 RNAi treatment. * indicates residual His-mycUb-independent pulldown of Ataxin-2-GFP, presumably by direct interaction with the Ni resin. D) Glossy eye phenotype caused by overexpression of Xbp1^spliced^ under the control of GMR-GAL4 (Genotype: eyFlp,GMR-GAL4,UAS-Xbp1s/FM7) E) Glossy eye phenotype is reduced in clones of suppressor 218. Genotype: eyFlp,GMR-GAL4,UAS-Xbp1s; FRT42D, Su218/FRT42D, ubiGFP. F) RNAi for Ataxin-2 suppresses the reduction of the “glossy” eye phenotype by clones of suppressor 218. Genotype: eyFlp,GMR-GAL4,UAS-Xbp1s; FRT42D, Su218/FRT42D, ubiGFP; UAS-Ataxin-2 RNAi.

To confirm the results obtained by mass spectrometry, we did immunoblots from biotin pulldowns of adult *Drosophila* head extracts expressing Ataxin-2-HA, ^bio^Ub and FLAG-HA-Fbxo42 in the eye (GMR-GAL4). Ubiquitylated Ataxin-2-HA was observed as 2 bands of slower mobility in the gel, with the top band (orange arrow), but not the lower band (blue arrow), being sensitive to the presence of DTT (Dithiothreitol) in the elution buffer (Fig. 2B), an indication that Fbxo42 could be promoting the ubiquitylation of Ataxin-2 in one or more cysteines (McClellan et al., 2019)(Kelsall, 2022)(Dikic and Schulman, 2023). Furthermore, in *Drosophila* S2 cells, we could detect Ataxin-2-GFP/His-myc-Ubiquitin conjugates that were diminished upon the presence of DTT in the elution buffer or RNAi depletion of Fbxo42 (Fig. 2C). Importantly, RNAi depletion of Ataxin-2 abolished the suppression of Xbp1^spliced^-induced glossy eye phenotype by Su218 (Fbxo42^N423*^) (Fig. 2D-F), indicating that Ataxin-2 is a relevant substrate of Fbxo42 in the context of the assay that was used in the original genetic screen (Fig.1). We confirmed the efficacy of the RNAi depletion of Fbxo42 in S2 cells (Fig. S2C) and Ataxin-2 in *Drosophila* eyes (Fig. S2D).

### Fbxo42 interacts with Ataxin-2

To investigate further the interaction between Fbxo42 and Ataxin-2, we did co-immunoprecipitation (co-IP) assays from S2 cells (Fig. 3A) and *Drosophila* heads (Fig. S3A) extracts. Ataxin-2-GFP can co-IP Fbxo42 in S2 cells (Fig. 3a) and FLAG-HA-Fbxo42 in *Drosophila* eyes (Fig. S3A). These results were confirmed by colocalization of Ataxin-2 and Fbxo42 in immunofluorescence assays. For these assays, we decided to use the ring gland, the larval organ that synthetizes ecdysone and other hormones that control molting throughout development (Mirth et al., 2005). The cells of the ring gland are large, have polytene chromosomes and have endogenous activation of Ire1 signaling, confirmed by the expression of Xbp1s-GFP (Fig. S3B), from a reporter of Ire1 activation, UAS-Xbp1-HA-GFP (Cairrão et al., 2022). Furthermore, ring gland cells express Ataxin-2 and Fbxo42 (Fig. 3B-E), and can easily be exposed to DTT, to induce high levels of ER stress and activation of the UPR. In untreated ring gland cells (Fig. 3B,C), Ataxin-2 and Fbxo42 have mostly uniform expression with little colocalization, although Fbxo42 presents higher levels of expression in the corpus allatum than in the prothoracic gland (Fig. 3B). In ring glands treated with 5 mM DTT for 4 hours (Fig. 3D,E), Ataxin-2 forms granules that are decorated with Fbxo42 foci (arrows in Fig. 3E and quantification in Fig. 3F). As an alternative method to induce the formation of Ataxin-2 granules in *Drosophila* cells, we used treatments with arsenite, as previously described (Buddika et al., 2020). Also in the case of arsenite treatments, we observed Fbxo42 foci decorating Ataxin-2 granules, both in the ring gland cells (Fig. S3C) and in the follicle cells of the ovary (Fig. S3D, E). Therefore, the recruitment of Fbxo42 to Ataxin-2 granules is not an exclusive of the induction of ER stress by DTT treatment.

**Figure 3.**
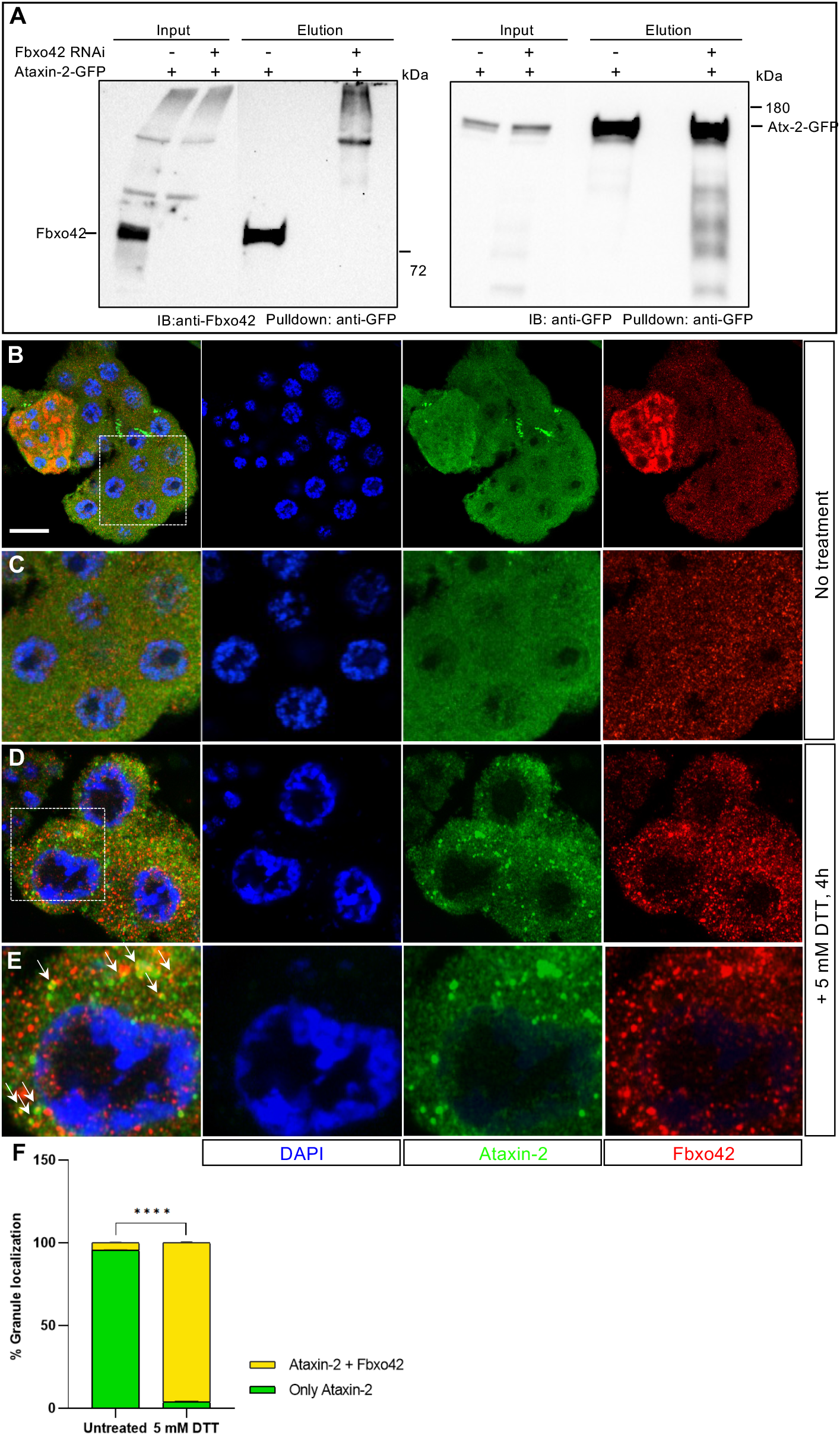
Fbxo42 interacts with Ataxin-2 in *Drosophila* tissues and S2 cells. A) Fbxo42 co-immunoprecipitates with Ataxin-2-GFP in S2 cells. Immunoblots probed with anti-Fbxo42 and anti-GFP antibodies from protein extracts of S2 cells expressing Ataxin-2-GFP before/ after immunoprecipitation with anti-GFP beads and with/without Fbxo-42 RNAi treatment. B) Immunofluorescence of ring gland (3^rd^ instar larva) showing uniform staining of Ataxin-2 (green) and Fbxo42 (red). DAPI is in blue. Scale bar = 30 μm. C) Inset of B). D) Immunofluorescence of ring gland (3^rd^ instar larva) after 4 h treatment with DTT (5 mM), to induce ER stress, shows Ataxin-2 (green) aggregates decorated with Fbxo42 (red). E) Inset of D). White arrows indicate examples of Ataxin-2 (green) aggregates decorated with Fbxo42 (red). F) Quantification of the number of granules containing Ataxin-2 only (green bar) or Fbxo42-decorated Ataxin-2 granules (yellow bar) present in untreated ring gland cells (shown in B) and in ring gland cells treated with 5 mM DTT for 4 h (shown in D). The quantification was done in 2 biological replicates per condition and is presented in percentage (%) as mean + SD. Two-way ANOVA coupled with Sidak’s multiple-comparison test, ^****^ *p* < 0.0001. The number of granules scored in untreated ring glands was n=277 (replicate 1) and n=269 (replicate 2). The number of granules scored in ring gland cells treated with DTT was n=552 (replicate 1, from 15 cells) and n=523 (replicate 2, from 11 cells).

### Fbxo42 promotes the degradation of Ataxin-2

Often, ubiquitylated proteins are degraded by the proteosome (Ciechanover and Schwartz, 1998). To investigate whether ubiquitylation of Ataxin-2 by Fbxo42 promotes the degradation of Ataxin-2, we did cycloheximide (CHX) chase experiments in S2 cells. In these experiments, we probed the stability of Ataxin-2 by immunoblot after 0, 6 and 12 hours of treatment of cells with CHX, an inhibitor of translation. Treating cells with CHX and MG132, an inhibitor of the proteosome, led to a significant stabilization of Ataxin-2-GFP (Fig. S4A), an indication that Ataxin-2-GFP can be degraded by the proteosome. Next, we found that Fbxo42 overexpression promotes the degradation of Ataxin-2-GFP (Fig. 4A,B), while Fbxo42 RNAi depletion protects Ataxin-2-GFP from degradation (Fig. 4C,D). We confirmed these results by doing immunofluorescence for Ataxin-2 in *Drosophila* 3^rd^ instar larvae eye discs containing clones of cells homozygous for Su218 (Fbxo42^N423*^), labelled by the absence of ubiGFP (Fig. 4E). Increased immunoreactivity for Ataxin-2 was observed in Fbxo42 homozygous mutant cells (arrows in Fig. 4E), but only when the imaginal discs were treated with 5 mM DTT for 4 hours, to induce ER stress, and only in the cells in the edge of the eye imaginal disc, presumably those exposed to higher amounts of DTT in the tissue.

**Figure 4.**
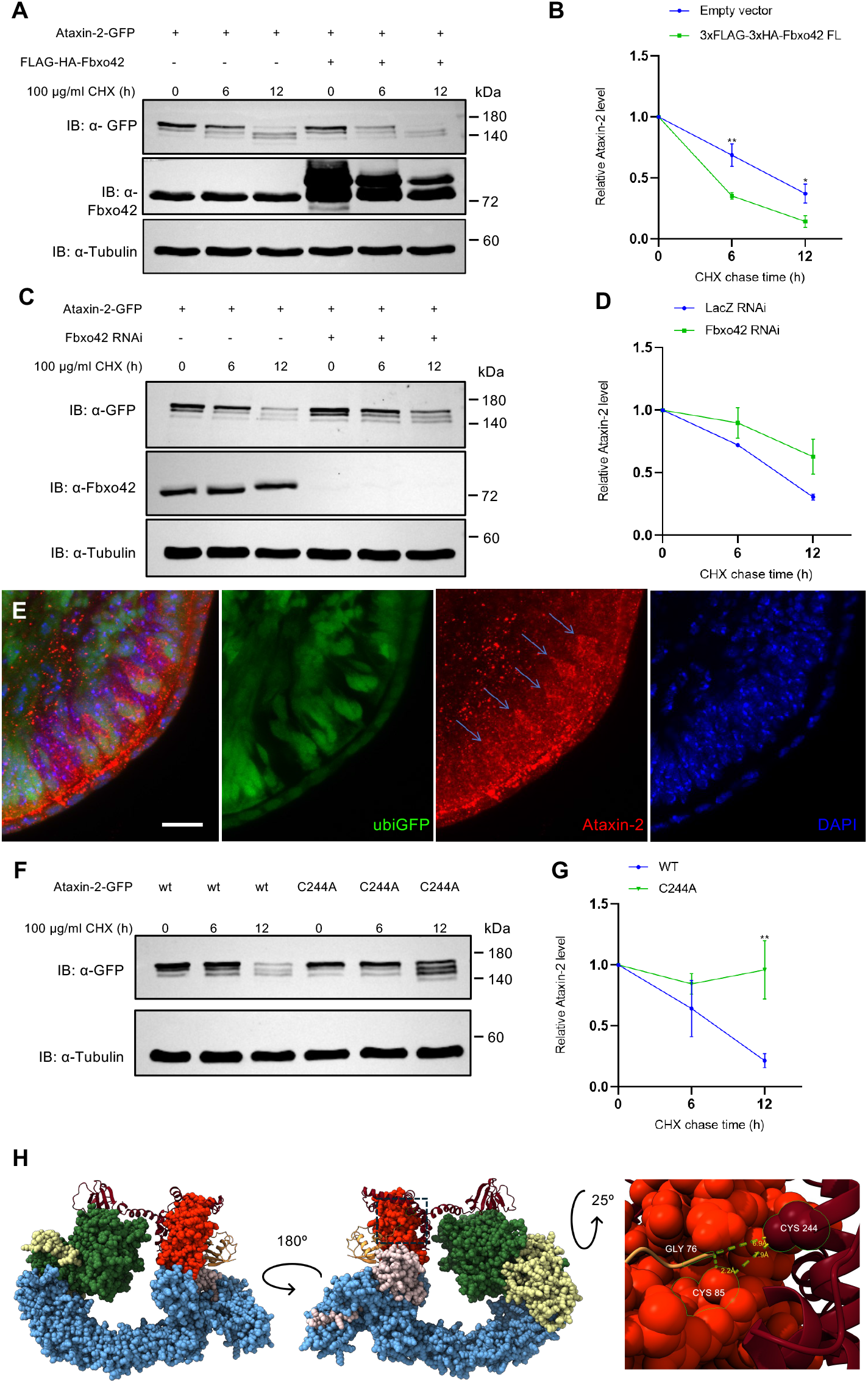
Fbxo42 promotes the degradation of Ataxin-2. A) Immunoblots probed with anti-Fbxo42 and anti-GFP antibodies from protein extracts of *Drosophila* S2 cells expressing Ataxin-2-GFP and Fbxo42 or Ataxin-2-GFP and empty vector negative control. S2 cells were treated with cycloheximide to inhibit protein translation and protein extracts were “chased” at the indicated time points. Tubulin is used as loading control. B) Quantification of Ataxin-2-GFP protein levels from A) and 2 other independent experiments is presented as mean + SEM. Two-way ANOVA coupled with Sidak’s multiple-comparison test, ^*^ *p* < 0.05 and ^**^ p < 0.01. C) Immunoblots probed with anti-Fbxo42 and anti-GFP antibodies from protein extracts of *Drosophila* S2 cells expressing Ataxin-2-GFP and Fbxo42 RNAi or Ataxin-2-GFP and LacZ RNAi, as negative control. S2 cells were treated with cycloheximide to inhibit protein translation and protein extracts were “chased” at the indicated time points. Tubulin is used as loading control. D) Quantification of Ataxin-2-GFP protein levels from B) and 2 other independent experiments is presented as mean + SEM. E) Immunofluorescence of 3^rd^ instar larva eye discs containing clones of Su218, labelled by the absence of ubiGFP (green). Endogenous ataxin-2 is in red and DAPI (blue) is a marker for nuclei. Whole brain/eye disc tissues were treated with 5 mM DTT (in PBS) for 4 hours, before fixation with formaldehyde. Genotype: eyFlp; FRT42D, Su218/FRT42D, ubiGFP. Scale bar = 10 μm. F) Immunoblots probed with anti-GFP from protein extracts of *Drosophila* S2 cells expressing Ataxin-2-GFP or Ataxin-2^C244A^-GFP. S2 cells were treated with cycloheximide to inhibit protein translation and protein extracts were “chased” at the indicated time points. Tubulin is used as loading control. G) Quantification of Ataxin-2-GFP (WT) and Ataxin-2^C244A^-GFP proteinlevels from f) and 2 other independent experiments is presented as mean + SEM. Two-way ANOVA coupled with Sidak’s multiple-comparison test, ^**^p < 0.01. H) Alphafold-3 prediction of a complex containing the LSM and LSM-AD domains (N57 to Q270) of Ataxin-2 (oxblood) together with Fbxo42 (green), Skp1 (yellow), Cullin1 (light blue), Rbx1 (pink), E2 (Effete, red) and Ubiquitin (orange). All proteins are represented as spheres except for Ataxin-2 and Ub that are in ribbon view. The right panel is a zoom in the region highlighted by the dashed rectangle, with a green highlight in Cys 244 of Ataxin-2, the catalytic Cys 85 of the E2 (effete) and the reactive C terminus of Ub (Gly 76).

To identify the cysteine residues in Ataxin-2 that are ubiquitylated upon overexpression of Fbxo42 we mutagenized C103, C154 or C244 to alanine (A) in Ataxin-2-GFP and did CHX chase experiments with these mutants, using S2 cells. While ataxin-2^C244A^ -GFP was stabilized (Fig. 4F,G), Ataxin-2^C103A^ -GFP and Ataxin-2^C154A^ -GFP (Fig. S4B) presented profiles of protein degradation similar to WT Ataxin-2-GFP. Interestingly, in an Alphafold-3 (Abramson et al., 2024) prediction displaying the LSM and LSM-AD domains (amino acids 62 to 269) of Ataxin-2 together with a Fbxo42/ Cullin-1/Skp-1/Rbx1/E2/Ub complex, the C244 of Ataxin-2 localizes in the vicinity of the reactive C terminus of Ub (Gly 76) and the catalytic Cys 85 of the E2 (Fig. 4H and S4C). Furthermore, Ataxin-2^C244A^-GFP did not present DTT-sensitive Ataxin-2-GFP/His-myc-Ubiquitin conjugates (Fig. S4D). We conclude that Fbxo42 promotes the degradation of Ataxin-2 by mediating the ubiquitylation of the C244 residue. Experiments with human cell lines have shown that loss of Fbxo42 leads to cell cycle progression delays and aberrant mitosis (Hundley et al., 2021), but the identity of the Fbxo42 substrates in this context remains unknown. Our results show that Fbxo42 is expressed in a variety of post-mitotic tissues, such as the ring gland or the adult *Drosophila* eye, and that Fbxo42 mediates the ubiquitylation and degradation of Ataxin-2 granules in these tissues.

### Ataxin-2 binds the Xbp1 mRNA during UPR activation

To investigate further the regulatory link between the RNA binding protein Ataxin-2 and Xbp1, we asked whether Ataxin-2 and Xbp1 mRNA could co-localize in cells undergoing activation of the UPR. We performed immunostaining with anti-Ataxin-2 and RNA fluorescent in situ hybridization (FISH) for Xbp1 mRNA in ring gland cells of *Drosophila* larvae treated with DTT, to induce high levels of UPR activation. In cells with no DTT treatment, Xbp1 mRNA is mostly detected in the cytoplasm, with little colocalization with Ataxin-2 (Fig. 5A). In DTT treated cells, Xbp1 mRNA colocalizes with cytoplasmic Ataxin-2 granules (arrows in Fig. 5B and quantification in Fig. 5C). Ataxin-2 is not present in Xbp1 mRNA nuclear foci (arrowheads in Fig. 5B) that presumably correspond to Xbp1 transcription, since it is known that ER stress and UPR activation activate Xbp1 transcription (Yoshida et al., 2001). To assess the specificity of our Xbp1 mRNA FISH probes, we used eye imaginal discs containing clones of Ex250 (Coelho et al., 2013)(Coelho et al., 2014), a deletion of most of *Xbp1*, that show strongly reduced FISH signal of Xbp1 mRNA in the Ex250 homozygous clones (Fig. S5A).

**Figure 5.**
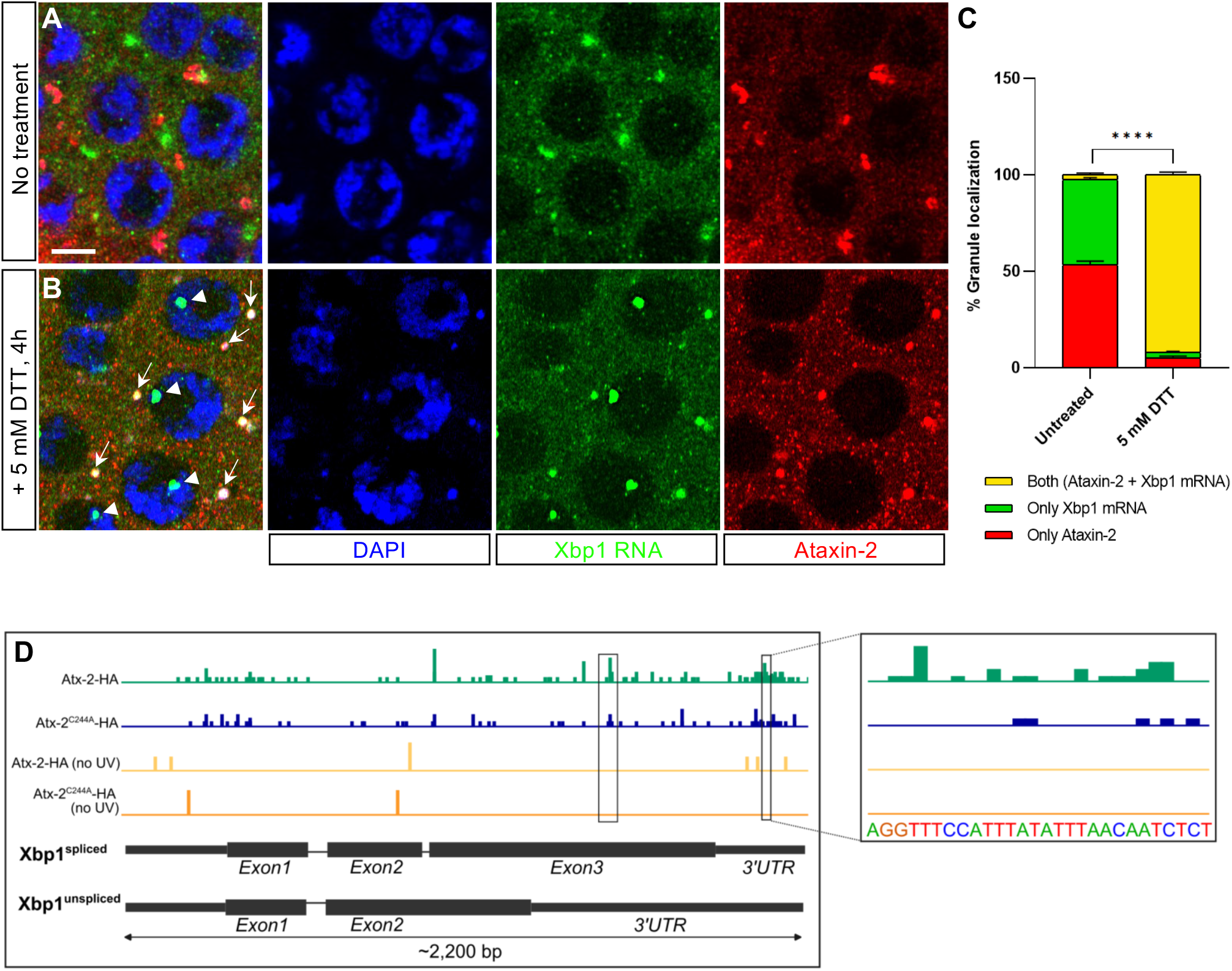
Ataxin-2 binds the Xbp1 mRNA during UPR activation. A) Immunofluorescence and RNA FISH of untreated ring gland cells (3^rd^ instar larva). Endogenous Ataxin-2 is in red (anti-Ataxin-2 antibody), Xbp1 mRNA in green (Stellaris FISH probes against Xbp1) and the nuclei is in blue (DAPI). Scale bar = 10 μm. B) Immunofluorescence and RNA FISH of ring gland cells treated with 5 mM DTT (to induce UPR activation) for 4h. Endogenous Ataxin-2 (red), Xbp1 mRNA (green) and nuclei (blue), as in a). Arrows indicate Xbp1 mRNA in Ataxin-2 granules. Arrowheads indicate Xbp1 mRNA in the nucleus. C) Quantification of the number of granules of Ataxin-2 protein only (green bar), Xbp1 mRNA only (red bar) or both Ataxin-2 protein and Xbp1 mRNA (yellow bar), in untreated ring gland cells (shown in A) and in ring gland cells treated with 5 mM DTT for 4 h (shown in B). The quantification was done in 2 biological replicates per condition and is presented in percentage (%) as mean + SD. Two-way ANOVA coupled with Sidak’s multiple-comparison test, ^****^ *p* < 0.0001. The number of granules scored in untreated ring glands was n=125 (replicate 1) and n=150 (replicate 2). The number of granules scored in ring gland cells treated with DTT was n= 123 (replicate 1) and n=156 (replicate 2). D) iCLIP results for Xbp1/Ataxin-2-HA. In green are depicted the peaks (sequencing reads) in *Xbp1* from UV-irradiated S2 cells transfected with Ataxin-2-HA. In blue are depicted the peaks in *Xbp1* of UV-irradiated cells transfected with Ataxin-2^C244A^-HA. The non-UV irradiated controls for Ataxin-2-HA and Ataxin-2^C244A^-HA samples are shown by the yellow and orange lines, respectively. To the right is a zoom of the *Xbp1* 3’UTR containing the AU-rich region.

These results suggest that, during UPR activation, Ataxin-2 binds to Xbp1 mRNA, which is found in cytoplasmic Ataxin-2 granules. To identify Ataxin-2 binding sites in the Xbp1 mRNA, we performed iCLIP (individual-nucleotide resolution UV crosslinking and immunoprecipitation) (König et al., 2010) experiments in S2 cells transfected with Ataxin-2-HA and treated with DTT, to induce high levels of UPR activation. Using an improved iCLIP protocol (Lee et al., 2021), we obtained results (Fig. 5D, S5B and S5C) showing that Ataxin-2 binds to the Xbp1 mRNA, especially to the 3’UTR, in a region containing an AU rich motif, which is consistent with the binding specificity previously identified for human and *Drosophila* Ataxin-2 (Yokoshi et al., 2014)(Singh et al., 2021). We also did iCLIP experiments with Ataxin-2^C244A^, the point mutation that protects Ataxin-2 from Fbxo42 mediated ubiquitylation and degradation. Interestingly, although Ataxin-2^C244A^ still binds Xbp1 mRNA, there is a reduction in the number of iCLIP crosslink reads, perhaps because C244 is in the LSM-AD domain that is important for RNA binding and essential to rescue the lethality of Ataxin-2 trans-heterozygous mutations (Bakthavachalu et al., 2018). Motif analysis around Ataxin-2 crosslink sites showed that various AU-rich pentamers (with a ‘C’ or ‘G’ or without) were enriched at the transcriptomic level in both wild-type and mutant Ataxin-2, but not in samples that were not UV-crosslinked (Fig. S5B). Moreover, the iCLIP crosslink positions (i.e. start positions of iCLIP cDNAs) were directly aligned with the motif centers at the metagene level (Figure S5C), including the GUUUC pentamer that was bound by Ataxin-2 on the Xbp1 3’UTR (Figure S5C).

Finally, we confirmed our findings in human (HeLa) cells (Figure S5D-F), where we also find an accumulation of Xbp1 mRNA in ataxin-2 granules in cells exposed to thapsigargin (1 μM) during 3 hours, to activate the UPR.

### Ataxin-2 and Fbxo42 regulate Xbp1 mRNA stability and translation during UPR activation

To test whether Ataxin-2 regulates the stability of Xbp1 mRNA, we performed actinomycin D (ActD) chase experiments in S2 cells, using the Xbp1 reporter for Ire1 activation, where the Xbp1^spliced^ form is tagged by GFP (Ryoo et al., 2007)(Cairrão et al., 2022). After ER stress induction with DTT (for 4h) and transcription inhibition with actinomycin D, the decay of the Xbp1 reporter mRNA was analyzed over time in S2 cells treated either with Ataxin-2 RNAi or control LacZ RNAi. Upon Ataxin-2 depletion, the stability of the Xbp1 reporter mRNA is decreased in comparison with the LacZ RNAi control condition (Fig. 6A). This result was confirmed by immunoblotting experiments (Fig. 6B), showing that, upon Ataxin-2 depletion and after DTT treatment (4 or 8 hours), the levels of Xbp1^spliced^-GFP protein are decreased in comparison with the LacZ RNAi control condition. Taken together, these results show that Ataxin-2 promotes Xbp1 mRNA stabilization and more Xbp1^spliced^ protein expression during UPR activation, being consistent with the results that show that human Ataxin-2 (Atxn-2), in general, stabilizes its target mRNAs and increases the abundance of the corresponding proteins (Yokoshi et al., 2014). Finally, we did immunoblotting experiments (Fig. 6C), showing that, upon Fbxo42 depletion and after DTT treatment (4 or 8 hours), the levels of Xbp1^spliced^-GFP protein are decreased in comparison with the LacZ RNAi control condition. This result demonstrates that Fbxo42 mediated degradation of Ataxin-2 is required for the translation of Xbp1 mRNA, which is consistent with the results obtained from the genetic screes, where loss of function mutation in Fbxo42 led to the suppression of the glossy eye phenotype caused by overexpression of Xbp1s.

**Figure 6.**
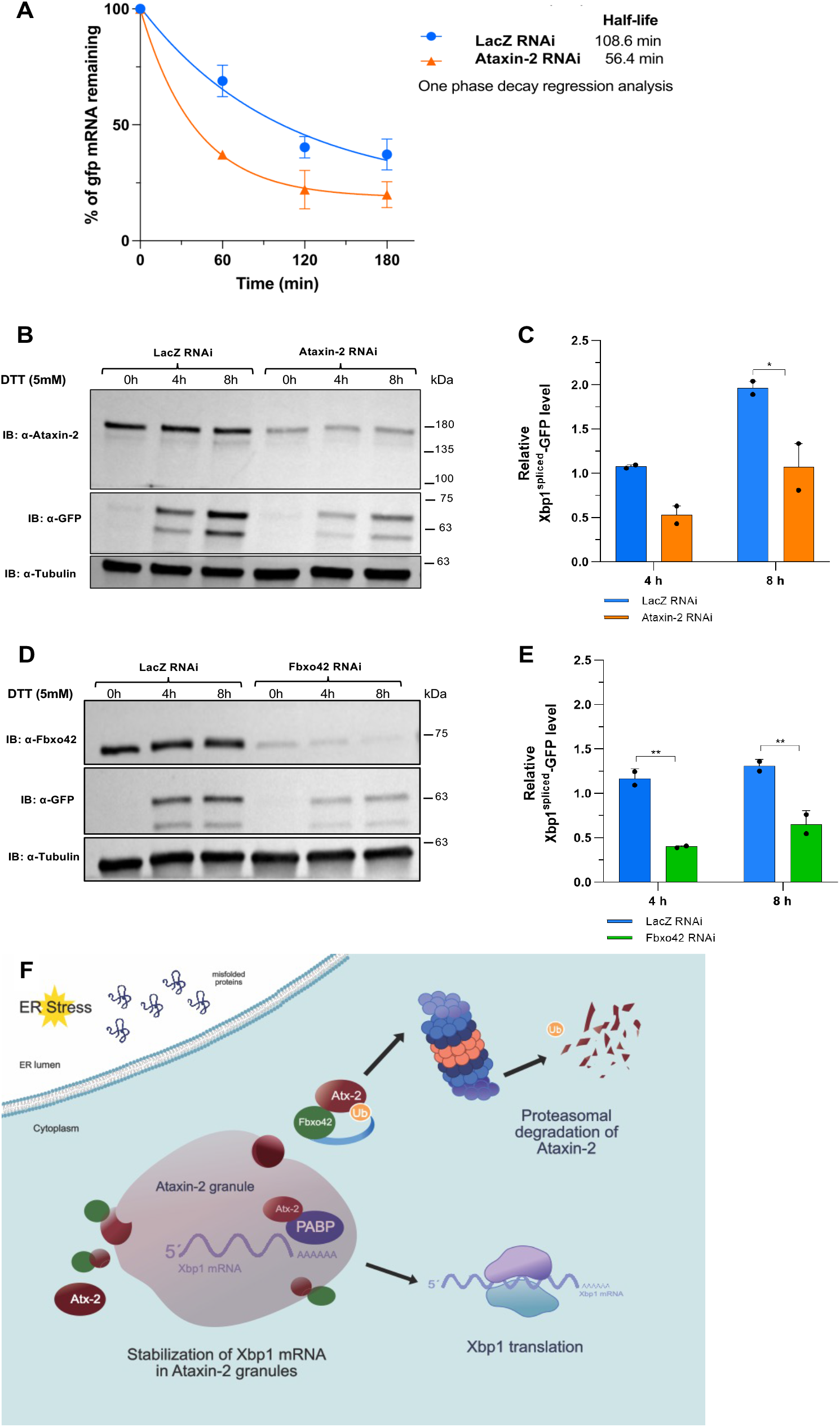
Ataxin-2 and Fbxo42 regulate the stability and translation of Xbp1 mRNA during UPR activation. A) Actinomycin D (ActD) chase experiments using S2 cells treated with Ataxin-2 RNAi or LacZ RNAi (control) and transfected with pUASTattb-Xbp1-HA-GFP. The cells were treated with 5 mM DTT (for 4 hours) and subsequently incubated with 5 µg/ml ActD until the indicated chase time points. B) Immunoblot of Xbp1^spliced^-GFP (probed with anti-GFP) and Ataxin-2 (probed with anti-Ataxin-2) after 5 mM DTT incubation for 4 and 8 hours. Tubulin was used as loading control. Protein extracts were harvested from cells treated with LacZ RNAi or Ataxin-2-RNAi and transfected with Xbp1-HA-GFP. C) Quantification of Xbp1^spliced^-GFP protein levels from B) and 1 other independent experiment is presented as mean + SD. Two-way ANOVA coupled with Sidak’s multiple-comparison test, *p < 0.05. D) Immunoblot of Xbp1^spliced^-GFP (probed with anti-GFP) and Fbxo42 (probed with anti-Ataxin-2) after 5 mM DTT incubation for 4 and 8 hours. Tubulin was used as loading control. Protein extracts were harvested from cells treated with LacZ RNAi or Fbxo42-RNAi and transfected with Xbp1-HA-GFP. E) Quantification of Xbp1^spliced^-GFP protein levels from D) and 1 other independent experiment is presented as mean + SD. Two-way ANOVA coupled with Sidak’s multiple-comparison test, ^**^p < 0.01. F) Upon ER stress, Xbp1 mRNA is transcribed and progressively accumulates in Ataxin-2 granules, where it is protected from degradation. Ataxin-2 (Atx-2) binds Xbp1 mRNA, together with Poly(A)-binding protein (PABP). Fbxo42 binds to Ataxin-2, promoting the ubiquitylation (Ub) of Ataxin-2 and its degradation by the proteosome. Xbp1 mRNA is released from Ataxin-2 and is translated.

## Discussion

In this study, we performed a genetic screen for suppressors of Xbp1^spliced^ induced “glossy” eye phenotype, leading to the identification of loss of function mutations in Fbxo42. Subsequently, we did a ^bio^Ub pulldown/proteomic experiment to identify Fbxo42 substrates, which led to the identification of Ataxin-2. Fbxo42 is recruited to Ataxin-2 granules during the activation of the UPR to ubiquitylate and promote the degradation of Ataxin-2. Finally, we show that Ataxin-2 binds to and stabilizes Xbp1 mRNA during the activation of the UPR, and that degradation of Ataxin-2 granules by Fbxo42 is required for the translation of Xbp1^spliced^.

We summarize our findings in the Fig. 6F. Upon high levels of ER stress, Xbp1 mRNA is spliced by the endoribonuclease activity of Ire1 and goes to Ataxin-2 granules, where it is protected from degradation but not translated. Polysome profiling experiments have shown that Ataxin-2 is found predominantly in non-ribosomal fractions (Yokoshi et al., 2014) and therefore, Ataxin-2-bound mRNAs are predominantly not translated. The progressive recruitment of Fbxo42 to Ataxin-2 granules leads to the ubiquitylation and degradation of Ataxin-2, releasing the Xbp1 mRNA free to be translated. In Fbxo42 mutant cells, Xbp1 mRNA remains not translated in Ataxin-2 granules, with less Xbp1 signaling readout (glossy eye phenotype), as in our initial genetic screen (Fig.1).

Our model is largely inspired by the results from the Ataxin-2/Fbxo42 immunofluorescence and Xbp1 mRNA FISH/Ataxin-2 immunofluorescence experiments (Figs. 3 and 5). Although, the Xbp1 mRNA FISH probes do not distinguish Xbp1^spliced^ from Xbp1^unspliced^, these experiments were done after 4 hours of exposure of the tissues to DTT, when most Xbp1 mRNA is already spliced by Ire1, but Xbp1s translation is still reduced (Majumder et al., 2012). In mouse embryonic fibroblasts, Xbp1s protein levels show a gradual increase between 3 and 9 hours of treatment with the ER stress inducer thapsigargin (Majumder et al., 2012), which is consistent with our studies in *Drosophila* S2 cells, where we typically see the highest levels of Xbp1s-GFP protein only upon 8 hours of DTT treatment (Fig. 6B,D and Cairrão et al., 2022). Our present results explain this temporal delay between the availability of Xbp1^spliced^ mRNA and the emergence of Xbp1^spliced^ protein – the Xbp1 mRNA stays untranslated in Ataxin-2 granules during the first few hours of ER stress and UPR activation.

Importantly, in ring gland cells, Ire1 signaling is activated even in the absence of DTT treatments (Fig. S3B), perhaps as part of their developmental role in the biosynthesis of the ecdysone hormone. However, the strong co-localization of Xbp1 mRNA with Ataxin-2 granules in ring gland cells only occurs upon DTT treatment and is concomitant with the strong transcriptional activation of Xbp1 (Fig. 5B). Therefore, regulation of Xbp1 mRNA by Ataxin-2 must be important when the UPR activation leads to high levels of Xbp1 transcription, for example by exposure of cells to DTT or overexpression of Xbp1^spliced^, as in our original genetic screen (Fig.1).

Cell death receptors (CD or DR) play a pivotal role in ER stress-induced cell death, including DR5 (Lu et al., 2014) and CD95 (Pelizzari-Raymundo et al., 2024). While the Ire1 RNase activity degrades the mRNAs of DR5 (Lu et al., 2014) and CD95 (Pelizzari-Raymundo et al., 2024) during an early, adaptative pro-survival stage (up to 6 hours of ER stress), the expression of DR5 and CD95 is upregulated at later stages, committing cells to the death fate. In this context, Xbp1s activates the transcription of CD95 to kill cells (Pelizzari-Raymundo et al., 2024). Therefore, while the mRNA of Xbp1 progressively accumulates in Ataxin-2 granules during the early, adaptative pro-survival stages, at later stages this large storage of Xbp1 mRNA could be quicky translated, upon the degradation of Ataxin-2 by Fbxo42, to trigger the transcriptional activation of cell death receptors and kill cells.

## Methods

### Plasmids

The cDNA clone GH02866 [Drosophila Genomics Resource Center (DGRC), DGRC Stock 6307; https://dgrc.bio.indiana.edu//stock/6307; RRID: DGRC_6307] was used to clone Fbxo42 into the Gateway® destination vectors pTFHW (to generate a N-terminally 3xFLAG-3xHA-tagged fusion protein) and pTWG (to create a C-terminally GFP-tagged fusion protein) following Invitrogen Gateway® protocol (Invitrogen). Primers used to generate pTFHW-Fbxo42 and pTWG-Fbxo42 are listed in table S2.

The pTFHW-ΔFbxo42 vector contains the deletion of the first 201 base pairs of Fbxo42 gene. This region encodes the first 67 amino acids of Fbxo42, and it includes the F-box domain. ΔFbxo42 sequence was cloned into the Gateway® destination vector pTFHW (primers in table S2).

The pUAST-Ataxin-2-GFP construct was generated by Gibson Assembly® (New England Biolabs, NEB). Briefly, the Ataxin-2 gene was amplified by PCR from flies expressing Ataxin-2 isoform A C-terminally fused with HA-tag (FlyORF F001031) and subcloned into the pJET1.2/blunt vector (CloneJET™ PCR Cloning Kit, Fermentas/Thermo Fisher Scientific), generating the pJET1.2-Ataxin-2-HA construct. Afterwards, Ataxin-2 sequence was PCR amplified from pJET1.2-Ataxin-2-HA and gfp was amplified from pUAST-Xbp1-EGFP (primers in table S2). Finally, the Ataxin-2 and GFP DNA fragments were cloned into the pUAST vector at XhoI (Fermentas/Thermo Fisher Scientific)/BglII (Thermo Fisher Scientific) restriction sites by Gibson reaction.

To generate the pUAST-Ataxin-2-HA construct, the DNA sequence encoding Ataxin-2 C-terminally fused with HA-tag was PCR amplified (primers in table S2) from pJET1.2-Ataxin-2-HA vector and cloned into pUAST plasmid at XhoI/BglII restriction sites by Gibson Assembly®. The Ataxin-2 mutations were generated by site-directed mutagenesis using the NZYMutagenesis kit (NZYtech) according to the manufacturer’s instructions. All constructs were confirmed by sequencing (Stabvida). The pUAST-His-Myc-Ubiquitin plasmid (Domingues and Ryoo, 2012) was kindly provided by HD Ryoo. The pMT-SkpA-HA and UAS-FLAG-Fblx7 (Bosch et al., 2014) plasmids were a gift from Iswar Hariharan.

### *Drosophila* stocks

Fly stocks and crosses were raised with standard cornmeal fly food, at 25°C under 12 h light/12 h dark cycles. New transgenic fly lines were made either at Bestgene or the Champalimaud Foundation transgenics facility. The fly stocks UAS-Xbp1u (Huang et al., 2017), UAS-Xbp1s (Ryoo et al., 2007), UAS-P35 (Grether et al., 1995), UAS-DIAP1 (Ryoo et al., 2002) and UAS-Rh1 (Kang and Ryoo, 2009) were gifts from Hyung Don Ryoo. The stock GMR-hid, eyFlp (Xu et al., 2005) was a gift from Andreas Bergmann. The stock UAS-FLAG-Fblx7 (Bosch et al., 2014) was a gift from Iswar Hariharan.

To identify the Fbxo42 substrates by mass spectrometry, flies bearing on the second chromosome a polyubiquitin chain conjugated with biotin and fused with E.coli BirA (biotin ligase), UAS-(bioUb)6-BirA (Ramirez et al., 2015a) (Franco et al., 2011), were crossed with flies carrying on the third chromosome the full-length Fbxo42 N-terminally tagged by 3xFLAG-3xHA-tag or the F-box control constructs: UAS-3xFLAG-3xHA-ΔFbxo42 (containing the deletion of the F-box domain) or UAS-FLAG-Fbxl7 (carrying a different fly F-box protein). Flies containing UAS-3xFLAG-3xHA-Fbxo42 without UAS-(bioUb)6-BirA were used as an additional control. The constructs were expressed in the Drosophila eye using the Glass Multimer Reporter-GAL4 driver (GMR-GAL4).

To confirm the mass spectrometry results, virgin female flies from the GMR-GAL4, UAS-(bioUb)6-BirA; UAS-FLAG-HA-Fbxo42 stock were crossed with male flies overexpressing Ataxin-2 isoform A C-terminally fused with 3xHA-tag (UAS-Ataxin-2-HA), obtained from FlyORF – Zurich ORFeome Project (fly line ID: F001031).

For the GFP pulldowns using fly heads, virgin female flies from GMR-GAL4, UAS-(bioUb)6-BirA; UAS-FLAG-HA-Fbxo42 or GMR-GAL4, UAS-(bioUb)6-BirA; UAS-FLAG-Fbxl7 stocks were crossed with male flies bearing UAS-Ataxin-2-GFP construct on the second chromosome.

Mutant clones in eye imaginal discs were generated by FLP (flippase recombinase)/FRT (FLP recombination target) recombination system (Golic, 1991), where flippase expression is under the control of the eyeless (ey) promoter (Newsome et al., 2000).

To express Ataxin-2 RNAi in Xbp1s-induced “glossy eye” with Fbxo42 mutant suppressor clones, female flies carrying one of the Fbxo42 suppressors, FRT42D, Su218 (Fbxo42N423*) were crossed with male flies bearing UAS-Ataxin-2 RNAi (BL 36114). Males from the progeny containing FRT42D, Su218 and UAS-Ataxin-2 RNAi were crossed with female flies with the genotype eyFlp, GMR-GAL4, UAS-Xbp1spliced; FRT42D, ubiGFP. Flies bearing eyFlp, GMR-GAL4, UAS-Xbp1spliced; FRT42D, Su218/ FRT42D, ubiGFP without UAS-Ataxin-2 RNAi were used as control.

### Mosaic genetic screen and mapping

The FRT42D stock was isogenized and males were collected for ethyl methanesulfonate (EMS) treatment. The males were starved for 8 hours in empty plastic vials and fed with a 25 mM EMS solution overnight for 16-18 hours (paper tissue soaked with EMS solution on the bottom of the vials). After EMS treatment, around 60 males were given one hour for recovery on tissue paper and were then crossed in mass to around 150 virgins of the genotype eyFlp, GMR-GAL4, UAS-Xbp1spliced; FRT42D, ubiGFP. The crosses were settled in new bottles every day, for four days in total and the F1 generation was screened for mosaic clones with suppression of the “glossy” eye phenotype. Suppressors of the “glossy” eye phenotype were balanced over CyO and retested for a reproducible phenotype. Only suppressor stocks that were homozygous lethal were kept, which was necessary for mapping of the mutations. For complementation analysis, each suppressor stock obtained from the screen was crossed to each other to bring the mutations in trans in the progeny to analyse whether unbalanced flies or flies only with the balancer CyO existed. When only CyO balanced progeny existed, the two suppressor mutations are lethal in trans and likely have a lethal hit in the same gene. For mapping of the suppressor mutations, we used the 2R deficiency kit (from the Bloomington Drosophila Stock Center). Virgins of the suppressor mutations balanced over CyO were crossed to males of each of the stocks bearing the balanced deficiencies and the F1 generation was screened for trans-heterozygosity. The existence of only balanced flies in F1 generation, indicated that the mutation is located within the region of the deficiency. The process was repeated with other smaller deficiencies until it could be defined the smallest region harboring the suppressor mutation. When available, we then tested lethal mutations in candidate genes in the region for trans-lethality with our mutations. The molecular identification of the mutations was done by genomic DNA sequencing of candidate genes from larvae homozygous for the suppressor mutations, collected from stocks balanced over CyO-GFP and also from the parental FRT42D stock before EMS mutagenesis. Genomic DNA preparation was done with the High Pure PCR Template Preparation kit from Roche and sequencing was performed by Stabvida.

### Anti-Fbxo42 antibody generation

To generate the anti-Fbxo42 rabbit polyclonal antibody, the DNA sequence coding for amino acids 384 to 667 of Fbxo42 was cloned into pETM-30 at XhoI/NcoI (NEB) restriction sites, to generate a N-terminal GST tagged protein. Beyond GST, Fbxo42 was tagged with two Histine (His)-tags placed on N- and C-terminals. Protein expression was performed in Rosetta bacterial cells. Supernatant and pellet extracts were run in 8% SDS-PAGE gels. Fbxo42 was purified by His-tag using a resin containing nickel [Profinity™ IMAC (Immobilized-Metal Affinity Chromatography) Ni-Charged Resin (cat. no. 156-0131, Bio-Rad)]. The purified Fbxo42 antigen in solution was sent to Eurogentec for antibody production, with subsequent in gel affinity purification of the antibody being performed in the laboratory. To minimize the background staining in immunofluorescence and immunoblot experiments, affinity purified anti-Fbxo42 antibody was preabsorbed with L1 larvae homozygous for Su226 (Fbxo42H436*).

### Immunofluorescence and imaging

*Drosophila* adult or larval tissues were dissected in 1× PBS, fixed with 4% PFA (paraformaldehyde) in 1× PBS at room temperature for at least 45 minutes and washed three times with PBT (1× PBS + 0.3% Triton X-100), 10 minutes each. Afterwards, fly’s tissues were incubated with primary antibodies diluted in BBT-250 (0.1% BSA, 0.1% Triton X-100, 250 mM NaCl in 1× PBS) overnight, at 4°C under gentle agitation. The primary antibodies used were as follow: rat anti-ELAV (1:200, 7E8A10, Developmental Studies Hybridoma Bank, DSHB), guinea-pig anti-Ataxin-2 (1:200, Zhang et al., 2013, a kind gift from Patrick Emery), mouse anti-HA (1:200, Covance, MMS101P) and rabbit anti-Fbxo42 (1:100). Incubation with anti-Fbxo42 antibody was preceded by an extra step of blocking and permeabilization; tissues were incubated in block-permeabilization solution (1% BSA, 0.3% Triton X-100 in 1× PBS) for 1 hour at room temperature. After overnight incubation with primary antibodies, fly’s tissues were washed three times with PBT for 10 minutes each and incubated with fluorescent conjugated secondary antibodies (Jackson ImmunoResearch Laboratories) for 2 hours at room temperature. Following three washes in PBT, the tissues were mounted in VECTASHIELD® Antifade Mounting Medium with DAPI (Vector Laboratories, H-1200-10) and image acquisition was performed on a confocal microscope (Leica SP5 Live or Zeiss LSM 880). For imaging of adult fly eyes, 2-5 days old flies were anesthetized with CO2 and transferred to a microscope slide where fly bodies were immobilized with transparent nail polish. Drosophila eye’ s pictures were acquired using a Leica Z16 APO macroscope equipped with a Leica DFC 420C camera with Leica’s extended depth of focus (Montage) software.

To promote UPR activation and the formation of Ataxin-2 granules, larval tissues were exposed to 5 mM DTT for 4 hours in PBS. As an alternative method to induce the formation of Ataxin-2 granules, larval tissues were treated with 0.5 mM arsenite for 2 hours. Untreated larval tissues were used as control.

Adult ovaries were dissected in 1× PBS and then transferred to Schneider’s *Drosophila* medium (Biowest). Ataxin-2 granules were induced by incubating ovaries with 0.5 mM arsenite in Schneider’s medium for 2 hours at 25°C (Gareau et al., 2013). Untreated ovaries were kept in Schneider’s medium and used as control.

### bioUb pulldowns

For the biotin pulldowns were used the aforementioned fly stocks carrying GMR-GAL4, UAS-(bioUb)6-BirA together with one of the following F-box constructs: UAS-FLAG-HA-Fbxo42; UAS-FLAG-HA-ΔFbxo42 or UAS-FLAG-Fbxl7, the latter two used as controls. Flies bearing only UAS-FLAG-HA-Fbxo42 without bioUb or BirA were used as an additional control. Adult flies with 2-5 days old were collected and fragmented by several rounds of flash freezing in liquid nitrogen and vortexing. Afterwards, fly heads were separated from the remaining body parts using a set of sieves with a nominal cut-off of 850, 710 and 425 µm on dry ice. Biotin pull-downs from *Drosophila* heads were performed as previously described (Ramirez et al., 2015b). Briefly, 500 mg of fly heads from each genotype was homogenized in 2.9 ml of lysis buffer (8 M urea, 1% SDS, 1× PBS, 50 mM N-ethylmaleimide from Sigma, and a protease inhibitor cocktail from Roche). After the centrifugation of lysates at 16,000g at 4 ºC for 5 min, the supernatant was applied to a PD10 desalting column (GE Healthcare) previously equilibrated with binding buffer (3 M urea, 1 M NaCl, 0.25% SDS, 1× PBS and 50 mM N-ethylmaleimide). Eluates, except 50 μl kept as input fraction, were incubated with 250 μl of NeutrAvidin agarose beads (Thermo Scientific) for 40 min at room temperature and further 2 h and 20 min at 4 ºC with gentle rolling. The material bound to the beads was washed with the following washing buffers (WB) (buffer composition can be found below): twice with WB1, thrice with WB2, once with WB3, thrice with WB4, once again with WB1, once with WB5, and thrice with WB6. The ubiquitylated material, still bound to the beads, was then eluted by heating the beads at 95 ºC for 5 min in 125 μl of elution buffer (250 mM Tris-HCl, pH 7.5, 40% glycerol, 4% SDS, 0.2% bromophenol blue, and 100 mM DTT). Finally, samples were centrifuged at 16,000g at room temperature for 2 min in a Vivaclear Mini 0.8 μm PES-micro-centrifuge filter unit (Sartorious) to discard the NeutrAvidin resin. The composition of the washing buffers used for the biotin pulldown assays were: WB1, 8 M urea, 0.25 SDS, 1× PBS; WB2, 6 M guanidine-HCl, 1× PBS; WB3, 6.4 M urea, 1 M NaCl, 0.2% SDS, 1× PBS; WB4, 4 M urea, 1 M NaCl, 10% isopropanol, 10% ethanol, 0.2% SDS, 1× PBS; WB5, 8 M urea, 1% SDS, 1× PBS; WB6, 2% SDS, 1× PBS.

Before proceeding with mass spectrometry, inputs and eluted fractions were analysed by immunoblotting with mouse anti-FLAG (1:1000, M2 clone, cat. no. F1804, Sigma-Aldrich), goat anti-Biotin-HRP (HRP, Horseradish Peroxidase-conjugated,1:1000) cat. no. 7075, Cell Signaling Technology, CST) and mouse anti-alpha-Tubulin (1:500, cat. no. AA4.3, DSHB) antibodies.

### In-gel trypsin digestion and peptide extraction

Ubiquitylated material eluted from biotin pulldown assays was concentrated in a Vivaspin 500 centrifugal filter units (Sartorius) and resolved by SDS-PAGE in 4–12% Bolt Bis-Tris precast gels (Invitrogen). Proteins were visualized with GelCode blue stain reagent (Invitrogen) and gel lanes were cut into 4 slices to remove known endogenously biotinylated proteins and avidin monomers, as previously described (Ramírez et al., 2021). Selected slices were subjected to in-gel trypsin digestion according to Shevchenko et al. (1996) with minor modifications (Shevchenko et al., 1996). Proteins were reduced with DTT (10 mM in 50 mM NH4HCO3, 56 ^°^C, 45 min), alkylated with chloroacetamide (25 mM in 50 mM NH4HCO3, room temperature, 30 min, dark) and incubated with trypsin (12.5 ng/ml in 50mM NH4HCO3, 37 ^°^C, overnight). The supernatant was recovered, and peptides were extracted twice from the gel: first, with 25mM NH4HCO3 and acetonitrile and then with 0.1% trifluoracetic acid and acetonitrile. The recovered supernatants and the extracted peptides were pooled, dried in a SpeedVac (Thermo Fisher Scientific) and subsequently desalted with homemade C18 tips (3M Empore C18).

### Liquid Chromatography with tandem Mass Spectrometry (LC-MS/MS)

An EASY-nLC 1200 liquid chromatography system interfaced with a Q Exactive HF-X mass spectrometer (Thermo Scientific) via a nanospray flex ion source was employed for the mass spectrometric analyses. Peptides were loaded onto an Acclaim Pep-Map100 pre-column (75 μm × 2 cm, Thermo Scientific) connected to an Acclaim PepMap RSLC C18 (75 μm × 25 cm, Thermo Scientific) analytical column. Peptides were eluted from the column using the following gradient: 18 min from 2.4 to 24%, 2 min from 24 to 32% and 12 min at 80% of acetonitrile in 0.1% formic acid at a flow rate of 300 nl min-1. The mass spectrometer was operated in positive ion mode. Full MS scans were acquired from m/z 375 to 1800 with a resolution of 120,000 at m/z 200. The 10 most intense ions were fragmented by higher energy C-trap dissociation with normalized collision energy of 28. MS/MS spectra were recorded with a resolution of 15,000 at m/z 200. The maximum ion injection times were 100 ms and 120 ms, whereas AGC target values were 3 × 106 and 5 × 105 for survey and MS/MS scans, respectively. In order to avoid repeat sequencing of peptides, dynamic exclusion was applied for 12 s. Singly charged ions or ions with unassigned charge state were also excluded from MS/MS. Data were acquired using Xcalibur software (Thermo Scientific).

### MS data processing and bioinformatics

Acquired raw data files were processed with the MaxQuant (Cox and Mann, 2008) software (version 1.6.0.16) using the internal search engine Andromeda (Cox et al., 2011) and tested against the UniProt database filtered for Drosophila melanogaster entries (release 2017_11; 43,868 entries). Mass tolerance was set to 8 and 20 ppm at the MS and MS/MS level, respectively. Enzyme specificity was set to trypsin, allowing for a maximum of three missed cleavages. Match between runs option was enabled with 1.5 min match time window and 20 min alignment window to match identification across samples. The minimum peptide length was set to seven amino acids. The false discovery rate for peptides and proteins was set to 1%. Normalized spectral protein LFQ (Label Free Quantitation) intensities were calculated using the MaxLFQ algorithm (Cox et al., 2014).

MaxQuant output data were analyzed with the Perseus module (version 1.5.6.0) (Tyanova et al., 2016) in order to determine the proteins significantly enriched in each of the genotypes. During the procedure, contaminants, reverse hits, as well as proteins with no intensity were removed, as well as those proteins only identified by site and/or with no unique peptides. Protein abundance was determined by LFQ intensity. Missing intensity values were replaced with values from a normal distribution (width 0.3 and down shift 1.8), meant to simulate expression below the detection limit (Tyanova et al., 2016). Statistically significant changes in protein abundance were assessed by two-tailed Student’s t test.

### Immunoblots from bioUb pulldowns

In order to confirm the results obtained by mass spectrometry, bioUb pulldowns were carried out using flies bearing GMR-Gal4, UAS-(bioUb)6-BirA; UAS-FLAG-HA-Fbxo42/UAS-Ataxin-2-HA. All Biotin pulldown steps were identical to those described above, with the modification of the elution step. The elution of biotinylated Ataxin-2-HA from the beads was performed with 4× Laemmli buffer (250 mM Tris-HCl pH 6.8, 40% glycerol, 8% SDS, 0.02% bromophenol blue) without reducing agents. Afterwards, the eluted volume was divided into two: half of the volume was incubated with 100 mM of DTT and heated at 65°C for 20 minutes. Input and eluted samples (in the presence/absence of DTT) were loaded onto 7.5% Mini-Protean® TGX™ (Tris-Glycine eXtended) Precast Protein Gels (Bio-Rad) and immunoblots were performed with rat anti-HA (1:1000, 7C9, Chromotek) antibody.

### Cell culture, transfections and RNAi treatments

*Drosophila* S2 cells (Schneider, 1972) were cultured at 24°C without CO2 in Schneider’s *Drosophila* medium (Biowest) supplemented with 10% heat inactivated Fetal Bovine Serum (FBS, Biowest), 100 U/ml of penicillin plus 100 µg/ml of streptomycin (Thermo Fisher Scientific). The complete medium was filtered with 0,2 µm PES filter (VWR). S2 cells were transfected with the indicated plasmids using Effectene transfection reagent (QIAGEN) according to the manufacturer’s instructions.

RNA-mediated interference (RNAi) was performed as described in (Clemens et al., 2000). Primer pairs bearing a 5’ end T7 RNA polymerase binding site were used to PCR amplify specific sequences of the genes to be inhibited. PCR products (with approximately 500-600 bp in length) were then purified with NucleoSpin® Gel and PCR Clean-up kit (Macherey-Nagel, MN) or with NZYGelpure kit (NZYtech) and used as templates for dsRNA synthesis with T7 RiboMAX™ Express Large Scale RNA Production System (Promega). After the DNase treatment, dsRNAs were purified and concentrated with RNA Clean & Concentrator™-5 kit (Zymo Research).

### Ubiquitylation assays in *Drosophila* S2 cells

For the ubiquitylation assay, 5.0 x 105 S2 cells/well were seeded in 6-well plates and treated with 25-30 µg of Fbxo42 dsRNA for 8 days. S2 cells endogenously expressing Fbxo42, and cells treated with Fbxo42 dsRNA were transfected with 200 ng of each one of the following plasmids: pActin-Gal4, pUAST-Ataxin-2-GFP and pUAST-His-Myc-Ubiquitin. Empty pUAST was used instead of pUAST-His-Myc-Ubiquitin as ubiquitylation control. 3 days after transfection, cells were incubated with 50 µM of MG132 (Z-Leu-Leu-Leu-al, C2211, Merck) for 4 hours, to inhibit proteasome activity, and resuspended in lysis buffer (50 mM Tris-HCl pH 7.5, 150 mM NaCl, 1 mM EDTA, 0.5% Triton X-100) supplemented with protease inhibitors (cOmplete™, Mini, EDTA-free Protease Inhibitor Cocktail, Roche) and 0.7% of NEM. Cell extracts were harvested and incubated on ice for at least 30 minutes. To perform the pulldown of proteins conjugated with His-Ubiquitin, each cell lysate was incubated with 15 µl of a nickel-charged resin suspension (Profinity™ IMAC Ni-Charged Resin, cat. no. 156-0131, Bio-Rad) for 2 hours and 20 minutes at 4°C. Afterwards, resin was washed thrice in 1× PBS + 20mM imidazole. The elution of the ubiquitylated molecules from the resin was performed with elution buffer containing 1× PBS + 250 mM imidazole at room temperature. The eluted volume was divided into two: half of the volume was incubated with 100 mM of DTT and heated at 65°C for 20 minutes. Then, 4× Laemmli buffer (240 mM Tris-HCl pH 6.8, 8% SDS, 40% glycerol, 250 mM DTT, 0.04% bromophenol blue) was added to the samples (with/without DTT), and these were boiled for 5 minutes at 95 °C. Finally, samples were loaded onto 4-15% Mini-Protean TGX™ Precast Protein Gels (Bio-Rad) and immunoblots were performed with rat anti-GFP (1:1000, 3H9, Chromotek) and rat anti-Myc (1:2000, 9E1, Chromotek) antibodies.

In order to observe the effect of C244A mutation on Ataxin-2 ubiquitylation profile, the ubiquitylation assay was repeated, in this case, S2 cells were co-transfected with pActinGal4 and pUAST-Ataxin-2C244A-GFP plasmids. All ubiquitylation assay steps were the same as those described above.

### Co-Immunoprecipitation

For co-immunoprecipitation of Fbxo42 and SkpA, S2 cells were transfected with pMT-SkpA-HA, pActinGal4, UAS-FLAG-HA-Fbxo42 or UAS-FLAG-HA-ΔFbxo42. Three days after, to induce the expression of SkpA-HA, 500 μM CuSO4 was added to the culture media 24 h before harvesting the cells in lysis buffer (20 mM HEPES 7.5 pH, 5 mM KCl, 1 mM MgCl2, 0.1% NP-40) supplemented with protease inhibitors (cOmplete™, Mini, EDTA-free Protease Inhibitor Cocktail, Roche). The immunoprecipitation was done using Anti-FLAG M2 Affinity gel (A2220, Sigma).

For co-immunoprecipitation of Fbxo42 and Ataxin-2, S2 cells endogenously expressing Fbxo42 protein or treated with Fbxo42 dsRNA were transfected with pActinGal4 and pUAST-Ataxin-2-GFP plasmids. Three days after, cells were resuspended in lysis buffer (50 mM Tris-HCl pH 7.5, 150 mM NaCl, 1 mM EDTA, 0.5% Triton X-100) supplemented with protease inhibitors (cOmplete™, Mini, EDTA-free Protease Inhibitor Cocktail, Roche).

To produce fly head extracts for Co-IP of Fbxo42 and Ataxin-2, flies expressing: GMR-GAL4, UAS-(bioUb)6-BirA/UAS-Ataxin-2-GFP; UAS-FLAG-HA-Fbxo42 or GMR-GAL4, UAS-(bioUb)6-BirA/UAS-Ataxin-2-GFP; UAS-FLAG-Fbxl7 were used. Fly heads were collected and homogenized in lysis buffer (50 mM Tris-HCl pH 7.4, 150 mM NaCl, 1mM EDTA, 1% Triton X-100) supplemented with protease inhibitors (cOmplete™, Mini, EDTA-free Protease Inhibitor Cocktail, Roche) using glass beads (0.5 mm Glass Beads, Scientific Industries) in a bead beater homogenizer.

Equal amounts of protein lysates were incubated with GFP-Trap agarose beads (ChromoTek GFP-Trap® Agarose) with rotation end-over-end for 2 hours and 30 minutes at 4°C. After that, tubes were centrifuged at 2700g for 2 minutes at 4°C. Then, GFP-Trap beads were washed three times in ice cold dilution buffer (10 mM Tris-HCl pH 7.5, 150 mM NaCl, 0.5 mM EDTA) supplemented with protease inhibitors (cOmplete™, Mini, EDTA-free Protease Inhibitor Cocktail, Roche). After the last wash, supernatants were removed and beads were resuspended in SDS-sample buffer (250 mM Tris-HCl pH 7.5, 40% glycerol, 4% SDS and 0.2% bromophenol blue). Finally, beads were boiled for 10 minutes at 95°C to dissociate immunocomplexes from the GFP-Trap beads and eluted fractions were recovered after centrifugation for 2 minutes at room temperature. In the case of the fly samples, the elution volume was divided into two: to half of the volume was added 100 mM DTT and this was then heated at 65°C for 15 minutes. Regarding the cell eluates, it was added 100 mM DTT to the entire eluted volume. Samples (inputs and eluates) were further analysed by western blot with rat anti-GFP (1:1000, 3H9, Chromotek) and rabbit anti-Fbxo42 (1:1000) antibodies.

### Cycloheximide chase experiments

Drosophila S2 cells were transfected with pActinGal4, wild-type or mutant versions of pUAST-Ataxin-2-GFP in the presence/absence of FLAG-HA-Fbxo42. Three days after transfection, cells were treated with 100 µg/ml cycloheximide (cat. no. J66901.03, Alfa Aesar) for 0, 6 and 12 hours, to inhibit the protein translation. S2 cells treated with Fbxo42 RNAi or with control RNAi and transfected with pActinGal4 and pUAST-Ataxin-2-GFP were also incubated with cycloheximide as aforementioned. At the indicated timepoints, cells were resuspended in lysis buffer (50 mM Tris-HCl pH 7.5, 150 mM NaCl, 1 mM EDTA, 0.5% Triton X-100) supplemented with protease inhibitors (cOmplete™, Mini, EDTA-free Protease Inhibitor Cocktail, Roche). Total cell extracts were analysed by immunoblotting with rat anti-GFP (1:1000, 3H9, Chromotek) and rabbit anti-Fbxo42 (1:1000) antibodies. Tubulin, detected with mouse anti-alpha-Tubulin antibody (1:500, AA4.3, DSHB) was used as loading control. Where indicated, 50 µM MG132 (Z-Leu-Leu-Leu-al, C2211, Merck) was added to the S2 cells to inhibit the proteosome.

### Immunoblots

Fly extracts or cell lysates were prepared in the above-mentioned lysis buffers supplemented with protease inhibitors (cOmplete™, Mini, EDTA-free Protease Inhibitor Cocktail, Roche). Protein concentration was determined by DC™ Protein Assay Kit from Bio-Rad. Afterwards, samples were mixed with 4× SDS Laemmli buffer (240 mM Tris-HCl pH 6.8, 8% SDS, 40% glycerol, 250 mM DTT, 0.04% bromophenol blue) and boiled at 95°C for 5 minutes. Proteins were size separated by SDS-PAGE and transferred onto nitrocellulose membranes of 0.2 µm (Bio-Rad). Immunoblots were blocked with 5% non-fat milk in TBS-T containing 0.1% Tween-20 or in PBS-T containing 0.05-0.1% Tween-20 for 1 hour. After blocking, membranes were incubated overnight at 4°C with the following primary antibodies (table S3): mouse anti-alpha-Tubulin (1:500, AA4.3, DSHB), rabbit anti-Fbxo42 (1:1000), rat anti-GFP (1:1000, 3H9, Chromotek), rat anti-HA (1:1000, 7C9, Chromotek), mouse anti-HA (1:1000, 16B12 clone, BioLegend), rat anti-Myc (1:2000, 9E1, Chromotek), guinea-pig anti-Ataxin-2 (1:5000) (kind gift from Patrick Emery), goat anti-Biotin-HRP (HRP, Horseradish Peroxidase) conjugated (1:1000, cat. no. 7075, CST) and mouse anti-FLAG (1:1000, M2 clone, cat. no. F1804, Sigma-Aldrich).

The following day, membranes were washed 4x for 10 minutes in TBS-T or PBS-T and incubated with HRP-conjugated secondary antibodies respective to the primary antibodies’ species, specifically, sheep anti-mouse HRP (1:10000, cat. no. NXA931, Cytiva), goat anti-rat HRP (1:2000, cat. no. R-05075-500, Advansta), goat anti-rabbit HRP (1:5000, product no. AS10 668, Agrisera) and goat anti-guinea pig HRP (1:5000, cat. no. PA128679, Thermo Scientific) for 2 hours at room temperature. Membranes were washed 4x for 10 minutes in TBS-T/PBS-T, and the signal was developed using ECL™ Prime Western Blotting Detection Reagent (RPN2232, Cytiva) and detected using a Chemidoc XRS+ (Bio-Rad) or an iBright™ FL 1500 Imaging System (Thermo Fisher Scientific). For Xbp1s-GFP immunoblot analysis, 3.0 x 106 cells were seeded in 25 cm2 flasks and treated with 30 µg of Ataxin-2 dsRNA or LacZ dsRNA (control) for 10 days. S2 cells were then re-plated in 12-well plates (4.5 x 105 cells/well) and transfected with pUASTattb-Xbp1-HA-GFP3’UTRspliced, pActinGal4 and empty pUAST plasmids. Three days after transfection, S2 cells were incubated with 5 mM DTT (to induce UPR activation) for 4 h and 8 h. Cell extracts were harvested at the indicated timepoints. Xbp1s-GFP protein levels were analyzed by immunoblot with rat anti-GFP (1:1000, 3H9, Chromotek) and tubulin was used as loading control, being detected with anti-alpha-Tubulin (1:500, AA4.3, DSHB). Ataxin-2 mRNA depletion was confirmed by RT-PCR before further analysis. The knockdown of Ataxin-2 protein was also analyzed by immunoblot with guinea pig anti-Ataxin-2 (1:5000). iBright™ Analysis Software was used for densitometry. Statistical analysis was performed using GraphPad Prism 8 software. Two-way ANOVAs were performed as mentioned in the figure legends. p values refer to: ^***^ p < 0.001, ^**^ p < 0.01 and *p< 0.05.

### AlphaFold 3 structural predictions

We utilized AlphaFold 3 (https://alphafoldserver.com/) to predict the structure of the Ataxin-2/Fbxo42/Cullin-1/Skp-1/ Rbx1/E2/Ub complex based on the protein sequences. Since AlphaFold struggles to accurately predict disordered or dynamic regions, we pre-processed the sequences by trimming large intrinsically disordered regions (IDRs), typically located in the N- and C-terminal tails. To identify these disordered regions, we employed the Metapredict V2 (Emenecker et al., 2021)(Emenecker et al., 2022) tool (https://metapredict.net/), which allowed us to remove these regions prior to structure prediction. This resulted in more stable poses with improved structural metrics. For the final AlphaFold 3 prediction, the sequences were: Ataxin-2 (aa 62 to 1084), Fbxo42 (aa 21 to 347 and 586 to 667), Cullin-1 (aa 11 to 774), Skp-1 (full length protein), Rbx1 (aa 47 to 122), E2(effete, full length protein) and Ub (full length protein). The resulting model was visualized using ChimeraX (Meng et al., 2023).

### Sequential Immunofluorescence and single molecule RNA fluorescence in situ hybridization (smFISH) in *Drosophila*

Single molecule RNA fluorescence in situ hybridization was performed with custom Stellaris RNA FISH probes labelled with Quasar 670 dye (LGC Biosearch Technologies) targeting the Xbp1 mRNA.

Sequential IF and smFISH was performed according to LGC Biosearch Technologies’ recommendations. Briefly, larvae were dissected in 1× PBS and fixed in 4% PFA in 1× PBS for 1 hour at room temperature. Tissues were then rinsed twice, 10 minutes each in 1× PBS followed by two washes in 0.1% Triton X-100. Subsequently, larval tissues were washed with 1× PBS and incubated with primary antibodies diluted in 1× PBS, overnight at 4°C with gentle agitation. Primary antibodies used were the following: mouse anti-ELAV (1:200) (cat. no. 9F8A9, DSHB) and guinea-pig anti-Ataxin-2 (1:200).

After overnight incubation with primary antibodies, tissues were washed three times with 1× PBS and incubated with appropriate secondary antibodies (Jackson ImmunoResearch Laboratories) diluted in 1× PBS for 2 hours at room temperature. Then, tissues were washed three times with 1× PBS, 10 minutes each, and incubated with freshly prepared Wash Buffer A, containing 1× Stellaris RNA FISH Wash Buffer A (cat. no. SMF-WA1-60, LGC Biosearch Technologies) and 10% (v/v) formamide (F9037, Sigma-Aldrich) in RNase free water, for 10 minutes at room temperature. Afterwards, samples were transferred to a 96-well plate and Wash Buffer A was discarded and replaced by freshly prepared Hybridization buffer [Stellaris RNA FISH Hybridization Buffer (cat. no. SMF-HB1-10, LGC Biosearch Technologies) + 10% (v/v) formamide] containing the Xbp1 probe set. Xbp1 probe stock (12.5 µM) was diluted 1:100 in Hybridization buffer to create a working probe solution of 125 nM. Larval tissues were incubated with Xbp1 probe solution in the dark at 37°C, overnight.

The following day, Xbp1 probe solution was discarded, and unbound probes were removed by incubation with Wash Buffer A in the dark at 37°C for 30 minutes. Tissues were mounted in VECTASHIELD® Antifade Mounting Medium with DAPI (Vector Laboratories, H-1200-10) and image acquisition was performed on a Zeiss LSM 880 confocal microscope.

Where indicated, the larval anterior parts’ (containing the ring glands and eye imaginal discs) were exposed to 5 mM DTT for 4 hours. Untreated tissues were used as control.

### Single-molecule fluorescence in situ hybridization and immunofluorescence (smFISH-IF) in HeLa cells and data analysis

The combined smFISH-IF experiments in HeLa cells were conducted as previously described (Gómez-Puerta et al., 2022). Briefly, HeLa 11ht cells in 1 mL culture medium (89% DMEM [Gibco], 10% (v/v) fetal bovine serum [Gibco] and 1% penicillin-streptomycin [Gibco]) were cultured in a 12-well plate (Greiner) at 37°C with 5% CO2. At 90% confluency, cells were either incubated with 1µM thapsigargin-treated (TG, Alomone) or untreated culture medium for 3 h. Cells were then washed twice with PBS and fixed with 4% paraformaldehyde (Electron Microscopy Sciences) for 10 min at room temperature (RT), followed by permeabilization with 0.2% Triton-X for another 10 min. Afterwards, a wash buffer (2 × SSC [Invitrogen] and 30% v/v deionized formamide [Ambion]) containing 3% (w/v) BSA (Sigma) was used to pre-block cells for 30min at RT. FISH probes targeting human XBP1 mRNA (Table S4) were designed using the anglerFISH probe designer and were made by enzymatic oligonucleotide labelling using Atto565-NHS as described before (Piskadlo et al., 2022). Cells were first hybridized with hybridization buffer (150 nM smFISH probes, 2 × SSC, 30% v/v formamide, 10% w/v dextran sulphate [Sigma]) for 4 h at 37 °C. Following three washes with wash buffer for 30 min, cells were incubated with primary antibody against human Ataxin-2 (Proteintech, 1776-1-AP, 1: 300) at RT for 4 h. After two washes with PBS containing 0.1% (v/v) Tween (Promega) for 30 min, the anti-rabbit IgG secondary antibody conjugated with ATTO 488 (Rockland, 611-152-122, 1: 5000) was added for incubation of another 30 min. Coverslips were washed twice with PBS before mounting them onto microscopy slides using ProLong Gold antifade reagent incl. DAPI (Molecular Probes).

smFISH-IF images were acquired on a Nikon spinning disk microscope equipped with a Yokogawa CSU W1 scan head, a Plan-APOCHROMAT 100 × 1.4 NA oil objective and iXon-888Life Back-illuminated EMCCD. Z-stacks were acquired with a step size of 0.2 μm. The maximum laser intensities were employed for all the channels, with exposure times of 1000 ms for Atto565 (200 gain), 1000ms for ATTO 488 (200 gain) and 200 ms for DAPI (400 gain), respectively.

Detection of single XBP1 mRNA spots in data from smFISH-IF experiments was performed in KNIME (Berthold et al., 2009) analogous to what was previously described (Voigt et al., 2019a, 2019b). Briefly, individual Z-slices were projected as maximum intensity projections. Following background-substraction, XBP1 mRNA spots were detected using threshold-based spot detection. Nuclear segmentation was performed on the DAPI signal using the Huang thresholding method while cytoplasmic segmentation was done using the background signal in the immunofluorescence (Ataxin-2) channel and a manual intensity threshold. Final masks were generated via substraction of nuclear from cytoplasmic segmentations and the number of XBP1 mRNAs per cell was quantified as the number of spots in each of these masks. To quantify XBP1-Ataxin-2 colocalization, mean Ataxin-2 IF intensities in all pixel positions obtained from XBP1 spot detection were quantified and normalized to the average Ataxin-2 background intensity of each cell. Results were exported for subsequent statistical analysis in Excel and GraphPad Prism 10.

### Individual nucleotide resolution UV cross-linking and immunoprecipitation (iCLIP)

The iCLIP method was carried out as described by Ule and collaborators (König et al., 2010) (Huppertz et al., 2014) with some modifications according to the improved iCLIP (iiCLIP) method (Lee et al., 2021), namely i) the introduction of an infrared adaptor as described in infrared-CLIP (irCLIP) (Zarnegar et al., 2016), ii) using reverse transcriptase (RT) Superscript IV and RT primers that contain carbon spacers as in irCLIP, and iii) ampure beads-based purification of cDNAs. Briefly, S2 cells transfected with Ataxin-2-HA or Ataxin-2C244A-HA were grown in 10 cm dishes. Three dishes were used per condition, including: a dish in which S2 cells were UV irradiated (+UV), a negative control in which no UV cross-linking was carried out (no UV) and an additional control in which UV irradiation was performed, although no antibody was used during the IP.

Three days after transfection, S2 cells of each condition were incubated with 5 mM DTT for 4 h (to induce UPR activation). After DTT incubation, the culture medium was removed, replaced by ice-cold PBS, and cells (from +UV and IgG dishes) were UV cross-linked twice with 0.15 J/cm2 in a Stratalinker 2400 (Analytic Jena) at 254 nm. Next, cells were harvested by scraping and the cell suspensions were centrifuged at 500g at 4°C for 5 min. Cell pellets were stored at - 80 °C until use.

Cell pellets were lysed in iCLIP lysis buffer (50 mM Tris-HCl pH 7.4, 100 mM NaCl, 1% Igepal, 0.1% SDS, 0.5% sodium deoxycholate) supplemented with cOmplete protease inhibitor cocktail (Roche), and the protein concentration was determined using a BCA Protein Assay Kit (Thermo Scientific). To digest the genomic DNA and obtain RNA fragments of an optimal size range, each cell lysate was incubated with Turbo DNase (Invitrogen) and 0.4 U/ml of RNase I (Thermo Scientific) for exactly 3 min at 37 °C with shaking at 1100 rpm. After incubation on ice for 5 min, lysates were spun at 4 °C at 13,000 rpm for 10 min, and supernatants were incubated with antibody-conjugated protein A/G Dynabeads (Thermofisher scientific) overnight at 4 °C. The antibodies were pre-conjugated to Protein A/G Dynabeads by incubating 100 µl of beads with 5 µg of either mouse Anti-HA (16B12 clone, BioLegend) or mouse IgG2b isotype control (MPC-11 clone, Tonbo Biosciences) for 1 h at room temperature. On the next day, supernatants were discarded, and beads were washed 3x with high-salt wash buffer (50 mM Tris-HCl pH 7.4, 1 M NaCl, 1 mM EDTA, 1% Igepal, 0.1% SDS, 0.5% sodium deoxycholate) and 1x with PNK wash buffer (20 mM Tris-HCl pH 7.4, 10 mM MgCl2, 0.2% Tween-20). Beads were then transferred to a new tube and the 3’ end dephosphorylation was carried out: the magnetic beads were resuspended in a mixture containing T4 Polynucleotide Kinase (Thermo Scientific) and FastAP alkaline phosphatase (Thermo Scientific). Samples were incubated for 40 min at 37 °C with shaking at 1100 rpm; beads were washed once with 1x ligation buffer (50 mM Tris-HCl pH 7.5, 10 mM MgCl2) and the on-bead ligation at room temperature (for 2 h) of the pre-adenylated adaptor L7-IR to the 3’ ends of the RNAs was performed. Samples were then washed thrice with high-salt wash buffer and once with PNK wash buffer. To remove the excess of adaptor, beads were resuspended in a removal mix containing 5’ deadenylase (NEB) and RecJf exonuclease (NEB) and incubated for 30 min at 30 °C followed by 30 min at 37 °C whilst shaking at 1100 rpm. Beads were washed 3x with high-salt wash buffer and 1x with PNK wash buffer and the protein-RNA complexes were eluted in 1x NuPAGE loading buffer + 100 mM DTT at 70 °C for 1 min. The protein-RNA complexes were separated by SDS-PAGE in a 4-12% NuPAGE Bis-Tris gel (Invitrogen), transferred to a nitrocellulose membrane (Cytiva) for 2 h (at 30V) and detected using infrared light on a Chemidoc MP (Bio-Rad).

The protein-RNA complexes were excised from the membrane using the infrared image printout as a mask, treated with a mix of proteinase K (Sigma-Aldrich) and PK+SDS buffer (10 mM Tris-HCl pH 7.4, 100 mM NaCl, 1 mM EDTA, 0.2% SDS) for 1 h at 50 °C whilst shaking at 1100 rpm. The membrane pieces were removed, samples were mixed with phenol: chloroform: isoamyl alcohol (Sigma-Aldrich), moved to a pre-spun 2 ml Phase Lock Gel Heavy tube and incubated for 5 min at 30 °C with shaking at 1100 rpm and phases were separated by centrifugation for 5 min at 13000 rpm. Next, chloroform (VWR) was added to the top phase, tubes were inverted to mix, and spun for 5 min at 13000 rpm at 4 °C. The aqueous layer was transferred into a new tube and the RNA was precipitated by adding 3 M sodium acetate pH 5.2 (Thermo Scientific), glycoblue (Invitrogen) and absolute ethanol, overnight at – 20 °C. Samples were spun for 30 min at 15000 rpm at 4 °C, the supernatants were discarded, and RNA pellets were washed with 80% ethanol and resuspended in DEPC-treated water.

The RNAs were reversely transcribed into cDNAs using the Superscript IV (SSIV, Thermo Scientific), dNTP mix and barcoded RT primers (IDT) according to the manufacturer’s instructions. Next, 1 M NaOH was added, and samples were incubated at 85 °C for 15 min for alkaline hydrolysis of the RNA. To neutralise the pH, the same amount of 1 M of HCl was added to the tubes. The cDNAs were purified using Mag-Bind Total Pure NGS beads (Omega Bio-Tek) following the manufacturer’s instructions. The cDNAs were eluted in DEPC-treated water, circularized in a mixture containing CircLigase II ssDNA ligase with betaine additive (LGC Biosearch Technologies) for 2 h at 60 °C, and purified using the Mag-Bind Total Pure NGS beads.

The cDNAs were PCR amplified using an optimal number of cycles in a Phusion HF master mix (Thermo Scientific) containing P5/P7 Illumina sequencing primers. The amplified cDNA library was gel purified, the 150 – 400 bp fragments were incubated in crush-soak gel buffer (500 mM NaCl, 1 mM EDTA, 0.05% SDS) at 65 °C for 2h 15 min, with 15 sec of agitation at 1100 rpm and 45 sec of rest, transferred to Costar SpinX columns (Corning), and spun at 13000 rpm for 5 min. The amplified cDNA was ethanol precipitated as described above, and resuspended in DEPC-treated water. Finally, the different (barcoded) cDNA libraries were mixed, purified using Mag-Bind Total Pure NGS beads, and sequenced at Genewiz (Azenta Life Sciences).

The iCLIP results were uploaded and analyzed on Flow bio (https://app.flow.bio) using the iCLIP pipeline as described https://docs.flow.bio/docs/clip-pipeline-1.0. In brief, 1) the demultiplexed reads were trimmed for the 3’end adapter sequence (AGATCGGAAGAGCACACGTCTGAA) by TrimGalore; 2) reads were pre-mapped to an index of non-coding RNA sequences using Bowtie; and the resulting unmapped reads mapped to the Drosophila genome (BDGP6.32 annotation) using STAR; 3) the aligned reads were deduplicated based on the unique molecular identifiers (UMI) and start position; 4) the crosslink positions were defined as the cDNA start position minus one in uniquely mapped reads; 5) positionally-enriched k-mer analysis (Kuret et al., 2022) as performed to determine enriched motifs in high-scoring crosslinks, using the following parameters: k-mer length as 5 nt (-k 5), proximal window as 20 nt (-w 20), distal window as 150 nt (-dw 150), and percentile parameter for assignment of ‘thresholded’ crosslinks to 0.7 (-p 0.7). All Ataxin-2 iCLIP data files generated, including the .bedgraph files of Fig. 5c, summary files of total crosslink counts in genes and RNA biotypes, and the parameters used in the iCLIP pipeline, are available on https://app.flow.bio/executions/562819420194487466. We selected the k-mers with highest PEKA scores and P value < 0.01 for Fig. S5B. For the alignment of crosslinks at motif centers in the 3’UTR (Fig. S5C), we normalized the crosslink counts at each meta position in the 3’UTR by the total number of crosslinks in the 3’UTR.

### mRNA stability assay

For mRNA stability experiments, 3.0 x 10^6^ S2 cells were seeded in 25 cm2 flasks and treated with 30 µg of Ataxin-2 dsRNA or LacZ dsRNA (control) for 10 days. S2 cells were then re-plated in 12-well plates (4.5 x 10^5^ cells/well) and transfected with pUASTattb-Xbp1-HA-GFP3’UTRspliced (Cairrão et al., 2022), pActinGal4 and pUAST plasmids. Three days after transfection, S2 cells were incubated with 5 mM DTT (to induce UPR activation) for 4 hours. Transcription was blocked with 5 µg/ml of actinomycin D (ActD, A9415, Sigma-Aldrich). Cells were incubated with ActD for 0, 1, 2 and 3 hours and samples were collected at each indicated timepoint for RNA extraction using NZYtech Total RNA Isolation kit. RNA samples were treated with Turbo DNase (Thermo) to eliminate DNA contamination. Equal amounts of total RNA were retro-transcribed using RevertAid H Minus First Strand cDNA Synthesis Kit (Thermo/Fermentas). Each qPCR reaction was performed on 1/40 of the cDNA obtained, using SSoFast EvaGreen Supermix (Bio-Rad) according to the manufacturer's instructions and Bio-Rad CFX-96 as detection system. For each sample, the levels of mRNAs were normalized using rp49 as a loading control. Normalized data were then used to quantify the relative levels of mRNA using the ΔΔCT method.

### Quantification and statistical analysis

The confocal images were analysed using the Fiji software and the ZEN Imaging Software (Zeiss). The number of granules and their components were quantified manually. Statistical analysis was carried out using GraphPad Prism 8 software. Two-way ANOVAs were performed as mentioned in the figure legends. p values refer to: ^****^ p < 0.0001, ^***^ p < 0.001, ^**^ p < 0.01 and ^*^ p < 0.05.

## Author Contributions

CCS designed the study, performed most experiments, analyzed data and wrote the manuscript. NS performed the genetic screen, identified the Fbxo42 mutants and made Fbxo42 constructs. FC designed and performed the mRNA stability/qPCRs and Fbxo42/SkpA co-IP experiments. JR, NO and UM performed mass spec identification of bioUB pulldowns and respective data analysis. VIR performed the genetic screen. CJG made Fbxo42 constructs. CA supervised and analyzed data. ZH supervised and analyzed FISH data. MY and FV performed FISH in HeLa cells. TNC performed the AF3 prediction. PG supervised and analyzed iCLIP data. PMD designed and supervised the study, performed the genetic screen, analyzed data and wrote the manuscript. All authors, except VIR, read and edited the manuscript.

## Acknowledgements

We would like to thank Hyung Don Ryoo, Andreas Bergmann, Iswar Hariharan, Patrick Emery, the Bloomington Drosophila Stock Center, FlyORF and the Developmental Studies Hybridoma Bank (DSHB) for fly stocks, plasmids and antibodies. We would like to thank Hyung Don Ryoo, Avi Ashkenazi and Brenda Schulman for comments on the manuscript. We thank Julia Domingos (Insta: cenasdajul) for the art work in Fig. 6F. The authors acknowledge Fundação para a Ciência e a Tecnologia, I.P. (FCT) for funding MOSTMICRO-ITQB, with references UIDB/04612/2020 and UIDP/04612/2020. The project was funded by the grants LCF/ PR/HR17/52150018 (‘la Caixa’ Foundation), FCT AGA-KHAN/ 541141368/2019 (FCT and Aga Khan Foundation), DL57/2016 to FC, SFRH/BD/130817/2017 to CCS and SFRH/BD/ 68361/2010 to NS. The proteomic analysis was performed in the Proteomics Core Facility-SGIKER at the University of the Basque Country (UPV/EHU).

---

**Figure S1.**
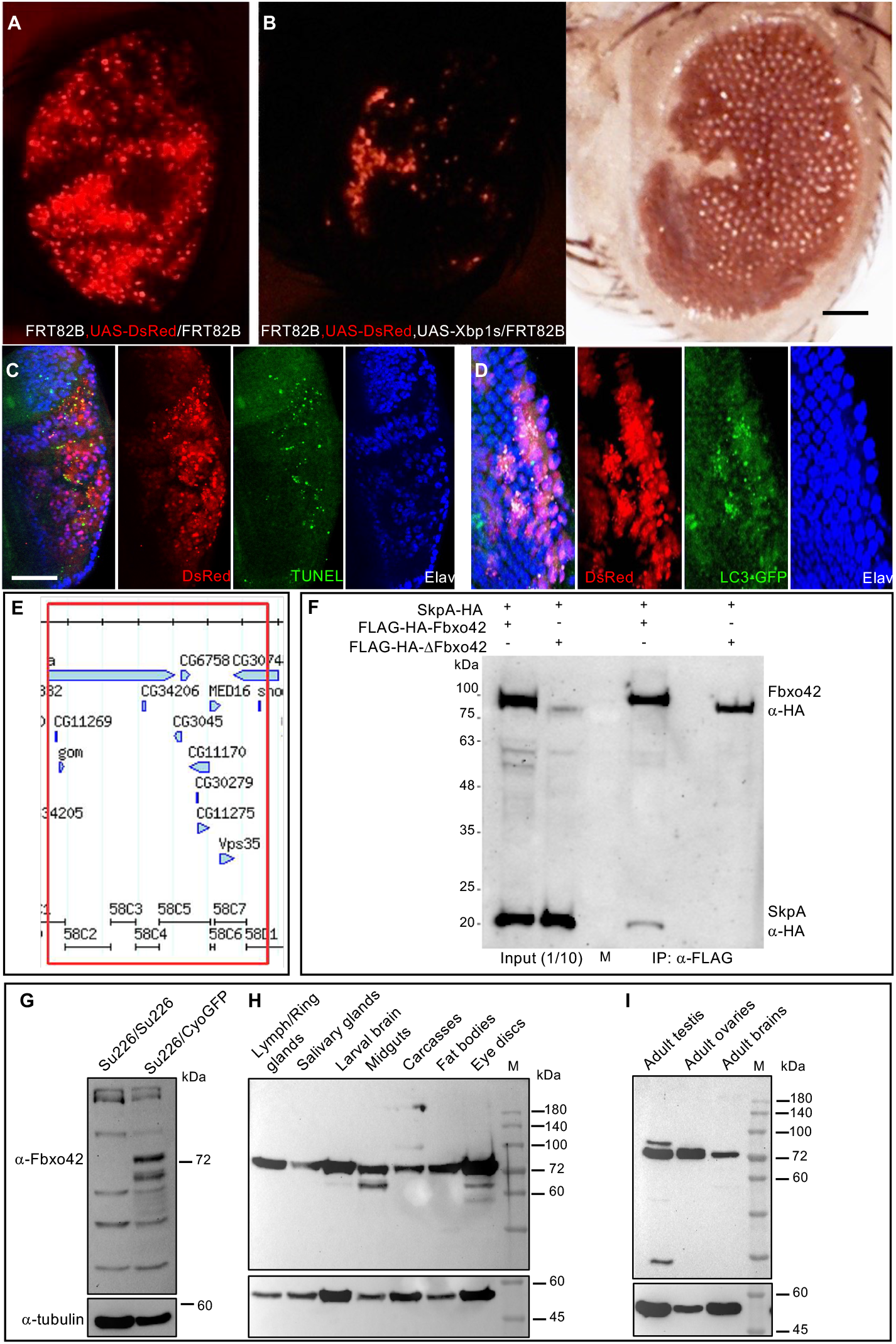
Overexpression of Xbp1^spliced^ activates apoptosis and autophagy in *Drosophila* larval eye discs and causes a glossy eye phenotype in adults. A) Adult *Drosophila* eye containing clones of cells (around 66% of the eye) with overexpression of DsRed. Genotype: eyFlp,GMR-GAL4; FRT82B, UAS-DsRed/FRT82B. B) Adult *Drosophila* eye containing clones of cells with overexpression of Xbp1s, labelled by DsRed, showing the glossy eye phenotype. Genotype: eyFlp,GMR-GAL4; FRT82B, UAS-DsRed, UAS-Xbp1s/FRT82B. Scale bar = 100 μm. C) Immunofluorescence of 3^rd^ instar larva eye discs containing clones of overexpression of Xbp1s, labelled by DsRed, showing increased TUNEL labelling (green), an apoptotic marker. ELAV (in blue) is a marker of the photoreceptors. Scale bar = 60 μm. D) Immunofluorescence of 3^rd^ instar larva eye discs containing clones of overexpression of Xbp1s, labelled by DsRed, showing increased LC3-GFP (green) labelling, a marker of activation of autophagy. ELAV (in blue) is a marker of the photoreceptors. E) Representation of the genomic region in 58C that was mapped by deficiencies, containing several candidates genes, including *Fbxo42 (CG6758)*. F) Immunoblot from S2 cell protein extract expressing SkpA-HA, FLAG-HA-Fbxo42 and FLAG-HA-ΔFbxo42 (a deletion of the Fbox domain that binds SkpA). SkpA-HA is co-immunoprecipitated by FLAG-HA-Fbxo42, but not by FLAG-HA-ΔFbxo42. G) Immunoblot from 1^st^ instar larva homozygous or heterozygous for Su226 (Fbxo42^H436*^) with an antibody raised against Fbxo42, showing the lack of the Fbxo42-specific band, just above the 72 kDa marker, in Su226 homozygous larvae. H) Immunoblot showing Fbxo42 expression in several from 3^rd^ instar larva dissected tissues. I) Immunoblot showing Fbxo42 expression in adult testis, ovaries and brain. Tubulin immunoblot serves as loading control in G-I.

**Figure S2.**
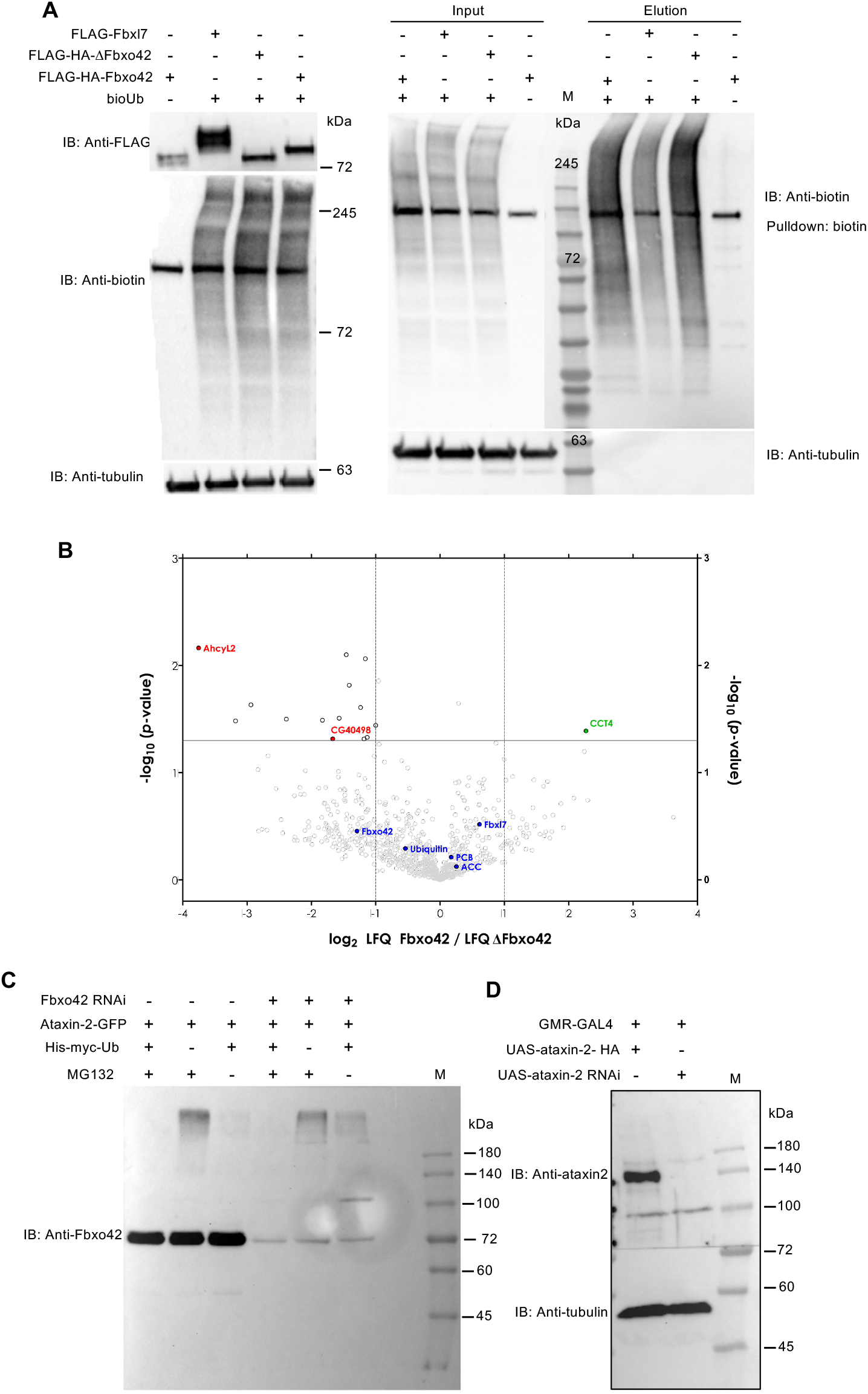
Fbxo42 promotes Ataxin-2/Ubiquitin conjugates in *Drosophila* eyes and S2 cells. A) Immunoblots of protein extracts before (input) and after (elution) streptavidin/biotin pulldowns from *Drosophila* adult heads expressing bioUb (ubiquitin with biotinylation acceptor site) and FLAG-HA tagged Fbxo42, ΔFbxo42 (deletion of the Fbox domain) or Fbxl7, under the control of GMR-GAL4. Biological triplicates were analyzed by mass spectrometry and the results are presented in volcano plots in Fig. 2A and Fig. S2B. B) Volcano plot of proteins identified by mass spectrometry after streptavidin/biotin pulldowns from *Drosophila* adult heads expressing ^bio^Ub (ubiquitin with biotinylation acceptor site) and Fbxo42 or ΔFbxo42, under the control of GMR-GAL4 driver. Results are presented as log2 LFQ (label free quantitation) intensity ratios. The color scheme of dots as in Fig 2a. Statistical analysis was performed by two-sided Student’s t test. The results are provided in table S1. C) Immunoblot of protein extracts from *Drosophila* S2 cells expressing His-myc-Ub, Ataxin-2-GFP and in the presence or absence of RNAi against Fbxo42 (as in Fig 2C) probed with an antibody against Fbxo42, attesting the efficiency of the RNAi treatments. D) Immunoblot of protein extracts from *Drosophila* heads expressing Ataxin-2-HA or Ataxin-2 RNAi, probed with an antibody against Ataxin-2, attesting the efficiency of Ataxin-2 RNAi treatments (as in Fig 2F).

**Figure S3.**
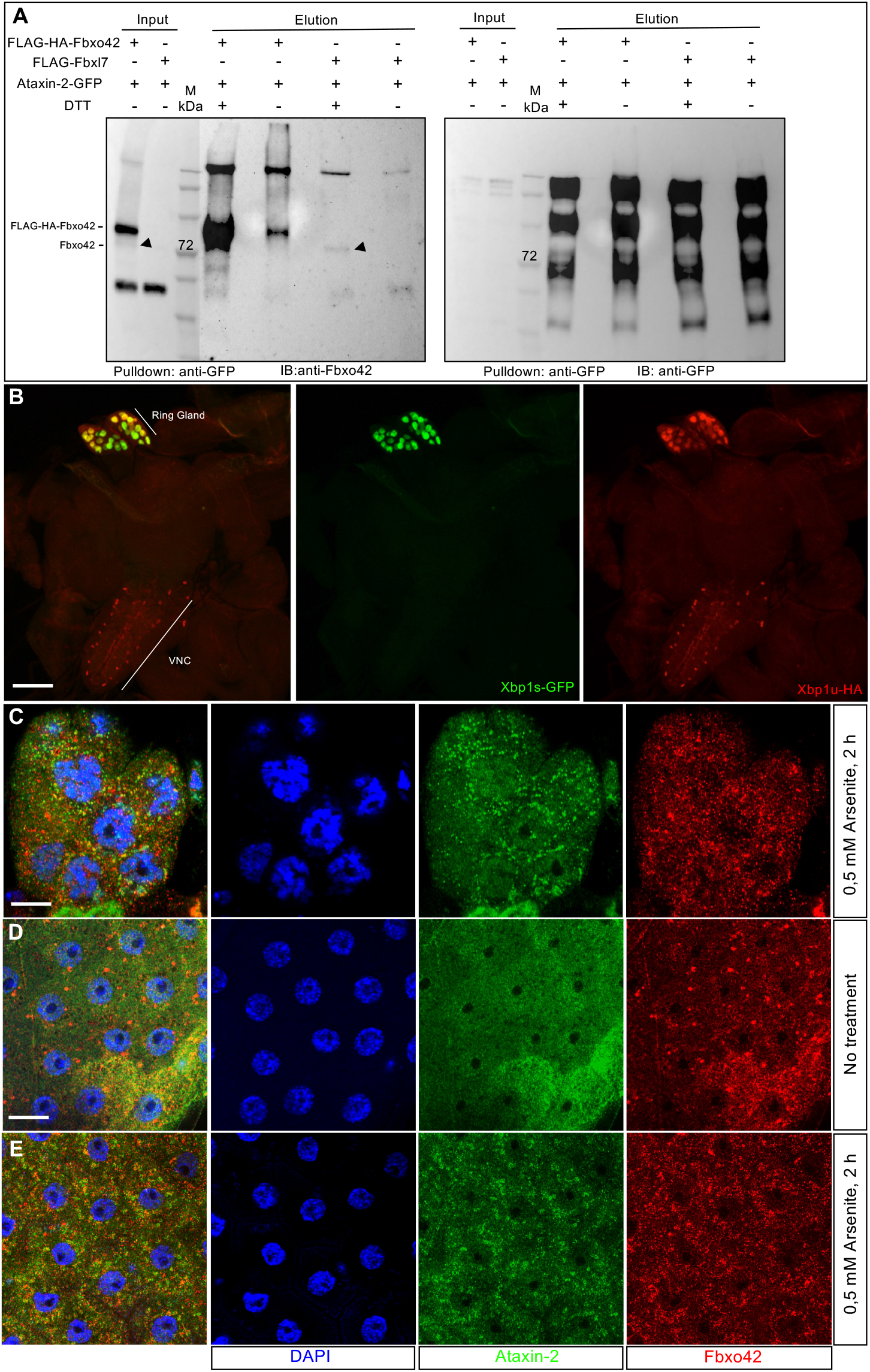
Fbxo42 interacts with Ataxin-2 in *Drosophila* eyes, ring glands and follicle cells of the ovary. A) Immunoblots probed with anti-Fbxo42 and anti-GFP antibodies from protein extracts of *Drosophila* adult heads expressing ataxin-2-GFP and Fbxo42 or Ataxin-2-GFP and Fbxl7 in the eye (under the control of GMR-GAL4 and all in the presence of ^bio^Ub), before/after immunoprecipitation with anti-GFP. Elution was done in the presence/absence of DTT. Black arrowhead indicates band with size corresponding to endogenous Fbxo42. B) Immunofluorescence of ring gland (3^rd^ instar larva) with attached brain/ventral nerve cord (VNC) expressing Xbp1-HA-GFP under the control of phm-GAL4, a ring gland driver. Expression of Xbp1^unspliced^-HA (red) is observed in the ring gland and in some cells in the VNC. Expression of Xbp1^spliced^-GFP (green) is observed in the ring gland, but not in the VNC. Scale bar = 100 μm. C) Immunofluorescence of ring gland (3^rd^ instar larva) after 2 h treatment with Arsenite (0,5 mM), shows Ataxin-2 (green) aggregates decorated with Fbxo42 (red). DAPI is in blue. Scale bar = 10 μm. D) Immunofluorescence of follicle cells of the adult ovary showing uniform staining of Ataxin-2 (green) and foci with increased Fbxo42 (red). DAPI is in blue. Scale bar = 10 μm. E) Immunofluorescence of follicle cells of the ovary after 2 h treatment with Arsenite (0,5 mM), showing Ataxin-2 (green) granules decorated with Fbxo42 (red). DAPI is in blue.

**Figure S4.**
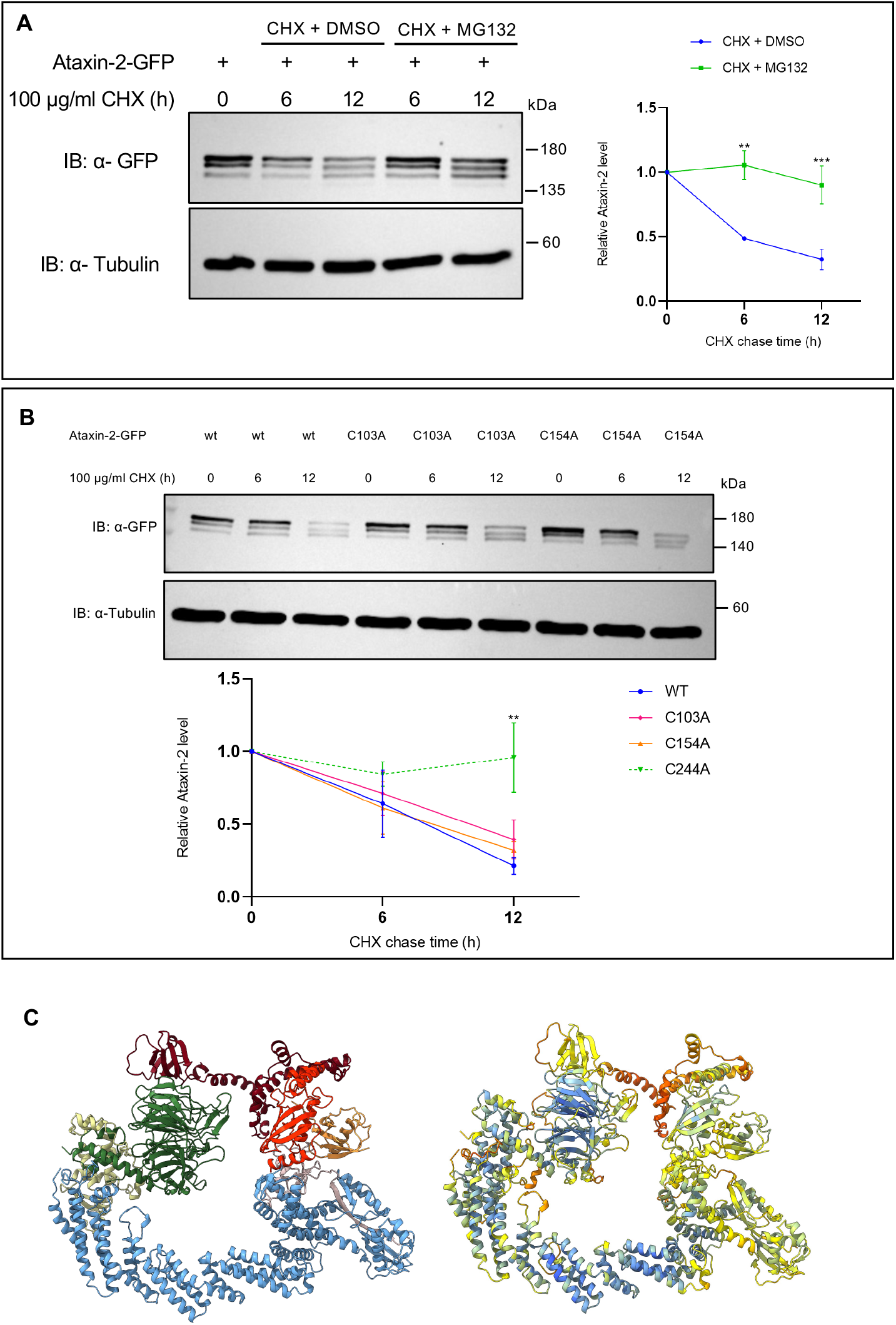
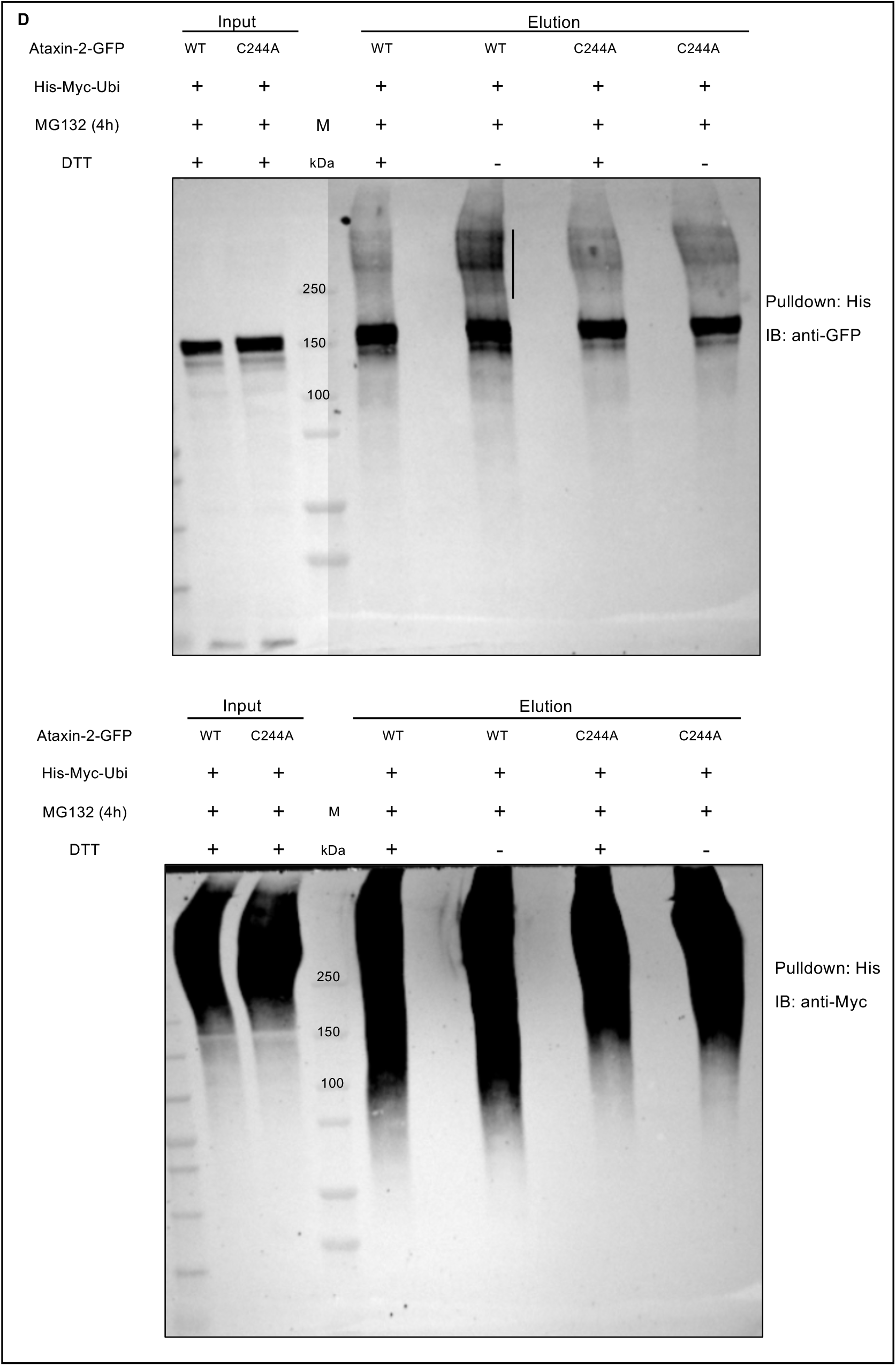
Fbxo42 promotes the degradation of Ataxin-2. A) Immunoblot probed with anti-GFP antibody from protein extracts of *Drosophila* S2 cells expressing Ataxin-2-GFP. S2 cells were treated with cycloheximide to inhibit protein translation and protein extracts were “chased” at the indicated time points, in the presence or absence of MG132, an inhibitor of the proteosome. Tubulin is used as loading control. Quantification of the protein levels of Ataxin-2-GFP (WT) from 3 independent experiments. Quantification for Ataxin-2-GFP is presented as mean + SEM. Two-way ANOVA coupled with Sidak’s multiple-comparison test, ^**^*p* < 0.01 and ^***^ p < 0.001. B) Immunoblots probed with anti-GFP from protein extracts of *Drosophila* S2 cells expressing Ataxin-2-GFP (WT, blue), Ataxin-2^C103A^ -GFP (pink) or Ataxin-2^C154A^ -GFP (orange). S2 cells were treated with cycloheximide to inhibit protein translation and protein extracts were “chased” at the indicated time points. Tubulin is used as loading control. Quantification of the protein levels of ataxin-2-GFP (WT) and C to A mutants from 3 independent experiments. Dashed green line represents the Ataxin-2^C244A^ -GFP from Fig. 4g. Quantification for Ataxin-2-GFP is presented as mean + SEM. Two-way ANOVA coupled with Sidak’s multiple-comparison test, ^**^ *p* < 0.01. C) Ribbon view of the Alphafold-3 prediction of a complex containing the LSM and LSM-AD domains (N57 to Q270) of Ataxin-2 (oxblood) together with Fbxo42 (green), Skp1 (yellow), Cullin1 (light blue), Rbx1 (pink), E2 (Effete, red) and Ubiquitin (orange), as in Fig. 4H, with the corresponding confidence values (pLDDT - Local Distance Difference Test). pLDDT values are color-coded on a scale from 0 to 100. Dark blue, pLDDT >90, light blue, 90>pLDDT>70, yellow, 70>pLDDT>50 and orange, pLDDT<50. D) Immunoblot of protein extracts from *Drosophila* S2 cells expressing His-myc-Ubiquitin (Ub) and Ataxin-2-GFP or Ataxin-2^C244A^ -GFP. S2 cells lysates were subjected to His pulldown and immunoblots with anti-GFP (top panels) or anti-myc (bottom panels). DTT sensitive His-myc-Ub conjugates (black vertical line) are detected for Ataxin-2-GFP but not for Ataxin-2^C244A^ -GFP.

**Figure S5.**
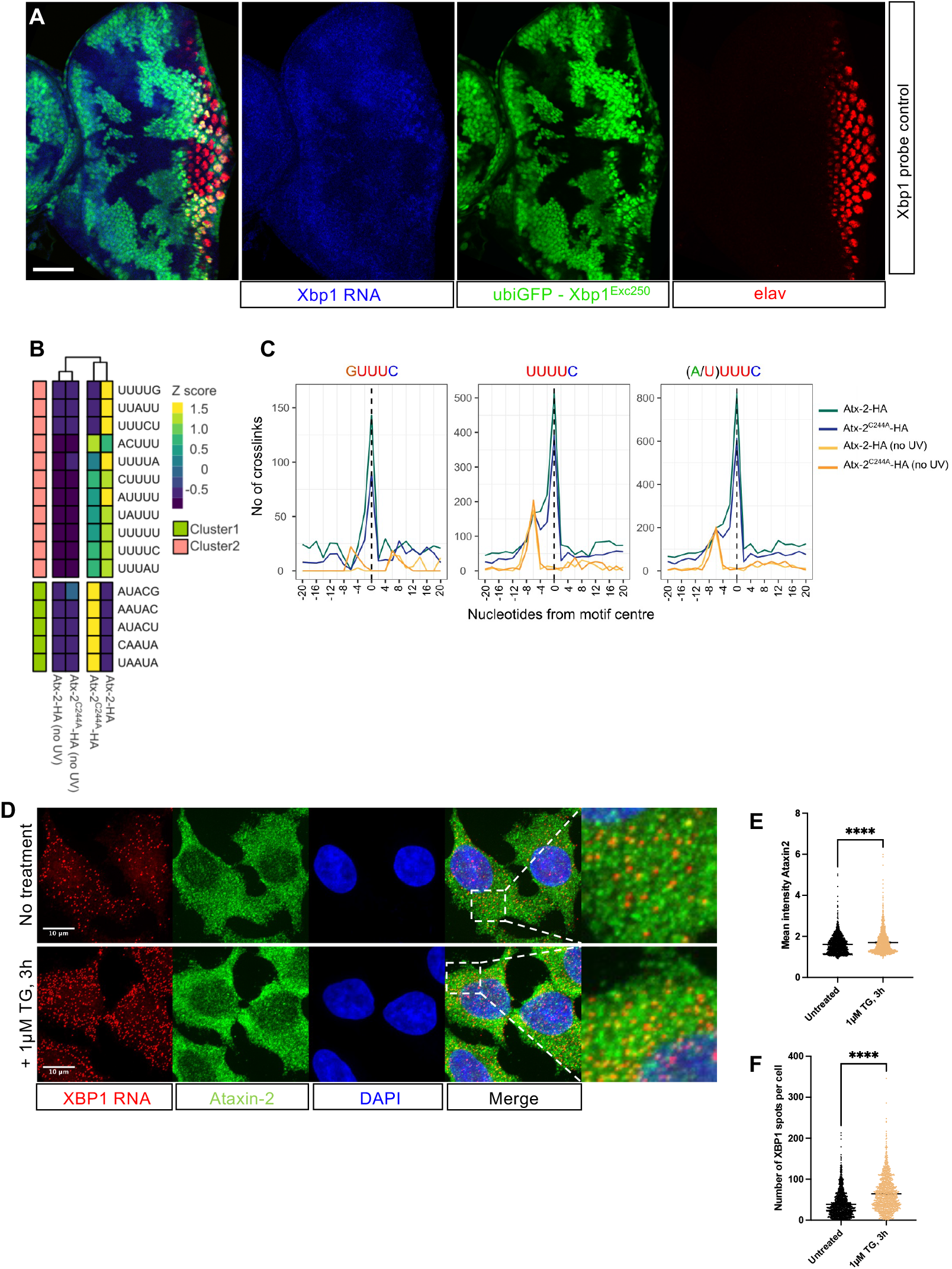
Xbp1 RNA-FISH probe control and additional iCLIP data. A) Immunofluorescence and RNA-FISH of 3^rd^ instar larva eye discs containing clones of cells homozygous for Xbp1^Exc250^, a deletion of Xbp1. Xbp1 mRNA levels, detected with Stellaris RNA FISH probes (in blue), are lower in Xbp1^Exc250^ homozygous cells, labelled by the absence of ubiGFP (green). The photoreceptors are labelled with ELAV (in red). Scale bar = 60 μm. B) Analysis of the motifs around the Ataxin-2-HA cross-link sites. The AU-rich pentamers identified in the transcripts of the UV-irradiated Ataxin-2-HA and Ataxin-2^C244A^-HA samples are labelled with different colors according to their Z-scores. C) iCLIP cross-link positions (i.e. start positions of iCLIP cDNAs) aligned with the motif centers at the metagene level. D) Combined single-molecule fluorescence in-situ hybridization (smFISH) and immunofluorescence (IF) analysis for colocalization of XBP1 mRNA (red) and Ataxin-2 (green) in fixed HeLa cells (DAPI = blue) with (or without) ER stress induced by 1µM thapsigargin (TG) for 3 hours. (E and F) Quantification of the data shown in (D). Depicted results were obtained from three biological replicates, incl. two technical replicates each. For each replicate and treatment, 10 images (app. 200 cells) were analyzed. E) Plot showing mean (per pixel) Ataxin-2 IF signal intensities across all cytoplasmic XBP1 mRNA spot positions (detected via smFISH) under control (untreated) and ER stress (1µM TG, 3h) conditions. Intensity values were normalized to the mean cytoplasmic background intensity per cell to correct for expression levels and ensure comparability in between replicates. Individual dots represent per-cell averages. Black bars show mean ± SEM. Statistical test: unpaired t-test, ^****^ : p-value <0.0001. F) Plot showing mean number of XBP1 mRNA spots per cell under the same conditions as in (E). Increased mRNA levels are indicative of XBP1 upregulation in response to UPR induction. Black bars show mean ± SEM. Statistical test: unpaired t-test, ^****^ : p-value <0.0001.

**Table S2:**
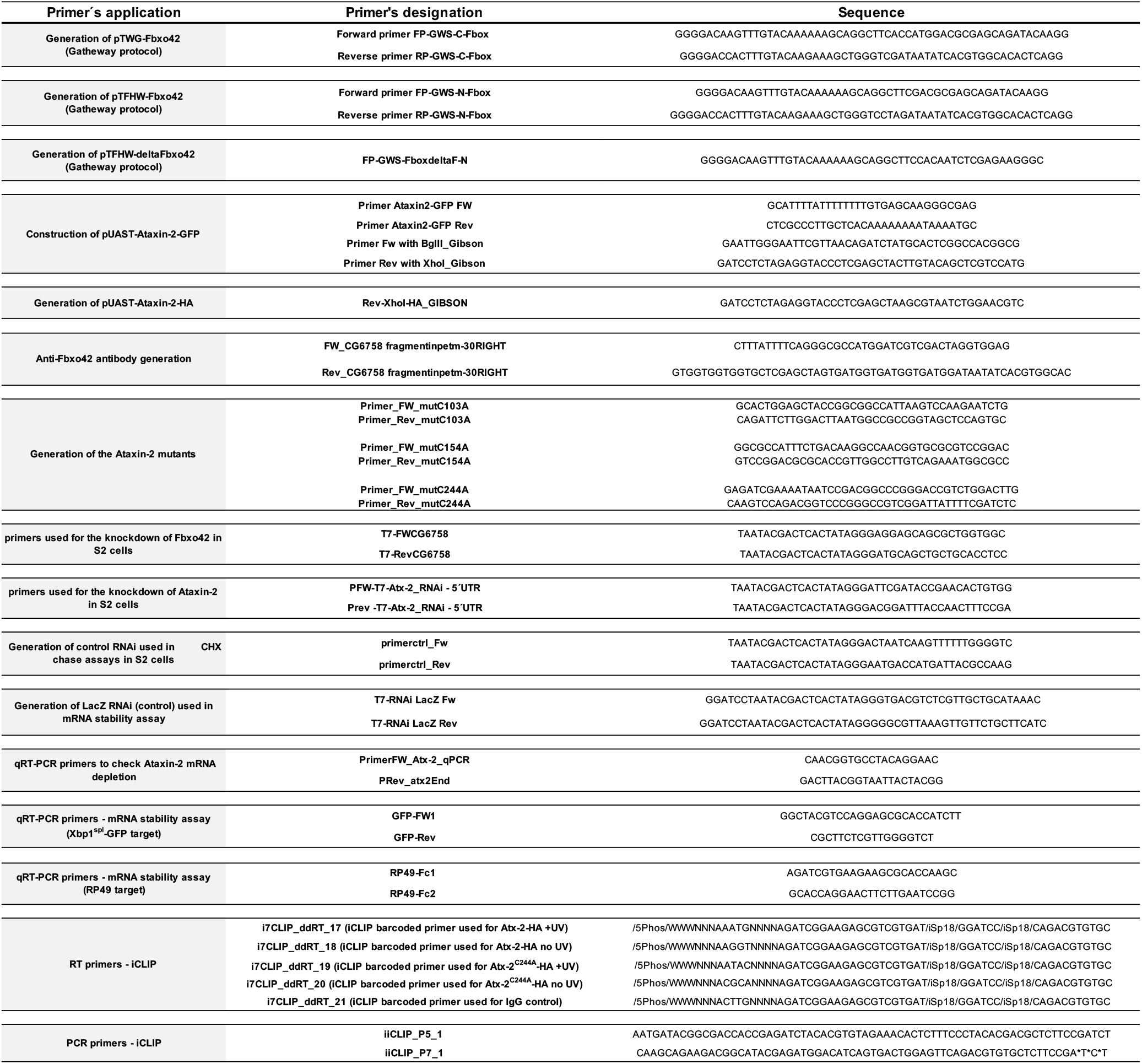
List of primers used in the present study.

**Table S3:**
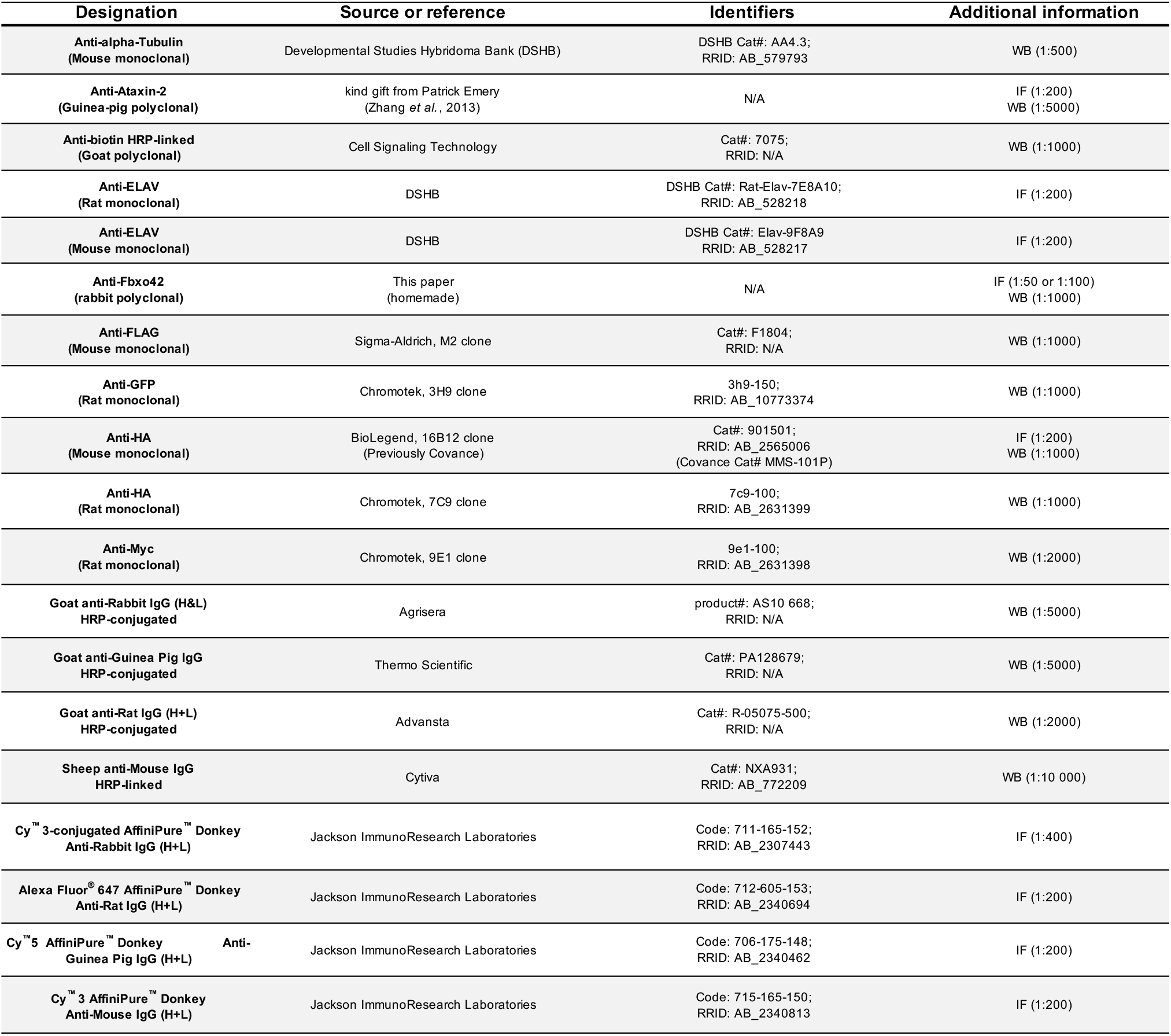
List of antibodies used in the present study.

**Table S4.**
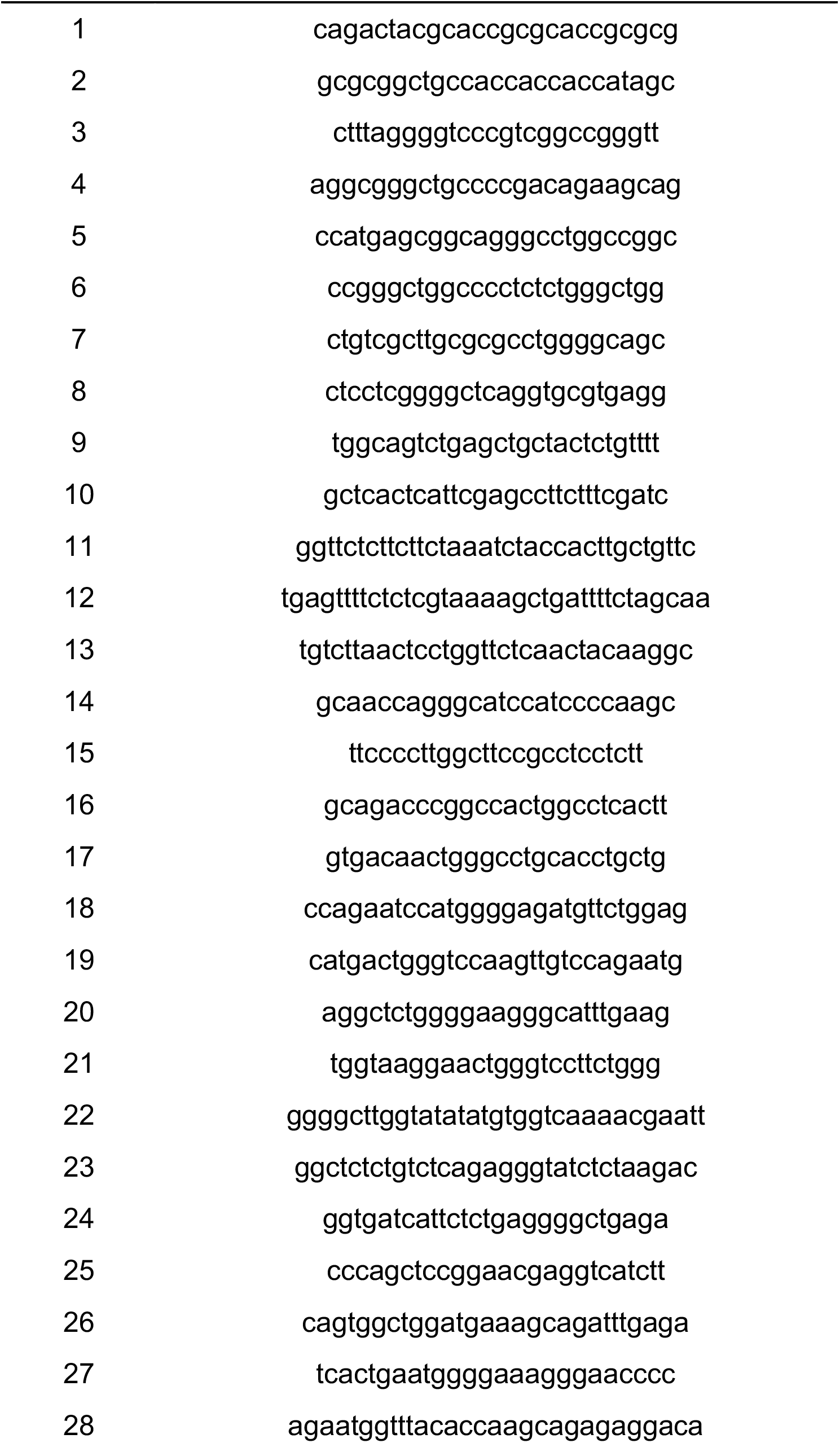
List of smFISH probes to detect human XBP1 mRNA.

